# Classification and Regression Models for Genomic Selection of Skewed Phenotypes: A Case for Disease Resistance in Winter Wheat (*Triticum aestivum* L.)

**DOI:** 10.1101/2021.12.16.472985

**Authors:** Lance F. Merrick, Dennis N. Lozada, Xianming Chen, Arron H. Carter

## Abstract

Most genomic prediction models are linear regression models that assume continuous and normally distributed phenotypes, but responses to diseases such as stripe rust (caused by *Puccinia striiformis* f. sp. tritici) are commonly recorded in ordinal scales and percentages. Disease severity (SEV) and infection type (IT) data in germplasm screening nurseries generally do not follow these assumptions. On this regard, researchers may ignore the lack of normality, transform the phenotypes, use generalized linear models, or use supervised learning algorithms and classification models with no restriction on the distribution of response variables, which are less sensitive when modeling ordinal scores. The goal of this research was to compare classification and regression genomic selection models for skewed phenotypes using stripe rust SEV and IT in winter wheat. We extensively compared both regression and classification prediction models using two training populations composed of breeding lines phenotyped in four years (2016-2018, and 2020) and a diversity panel phenotyped in four years (2013-2016). The prediction models used 19,861 genotyping-by-sequencing single-nucleotide polymorphism markers. Overall, square root transformed phenotypes using rrBLUP and support vector machine regression models displayed the highest combination of accuracy and relative efficiency across the regression and classification models. Further, a classification system based on support vector machine and ordinal Bayesian models with a 2-Class scale for SEV reached the highest class accuracy of 0.99. This study showed that breeders can use linear and non-parametric regression models within their own breeding lines over combined years to accurately predict skewed phenotypes.

## 1 Introduction

Genomic selection (GS) is posed to increase genetic gain and reduce cycle time for complex agronomic traits that are difficult to phenotype and analyze. With the advent of high-throughput genotyping, it is now feasible to develop and implement GS models for categorical/ordinal phenotypes that are common in most breeding programs and often difficult to analyze. The difficulty in phenotyping and analysis can be due to the traits’ genetic complexity, environmental dependency to display variation, and the inability of statistical models to model phenotypes adequately. Most GS models are linear regression models that assume continuous and normally distributed phenotypes (Montesinos-López et al., 2015c). When faced with data that do not follow the assumption of a linear model, researchers have several options. They may either ignore the lack of normality, transform the phenotypes, use generalized linear models (GLM) or use machine learning (ML) algorithms and classification models. Machine learning models have no restriction on the distribution of response variables which are less sensitive when modeling ordinal scores (Montesinos-López et al., 2015a; González-Camacho et al., 2018). Most GS models treat disease resistance as continuous values and utilize regression models and transformations for prediction whereas only a few studies have used classification methods (Ornella et al., 2012, 2014; Rutkoski et al., 2014; Muleta et al., 2017).

When the number of categories is large and the data follows more of a normal distribution, the ordinality of data can be ignored (Montesinos-López et al., 2015b). However, ignoring the lack of normality and using linear regression models imposes various problems. Linear regression models are limited to modeling additive effects only, whereas machine learning models account for both non-additive and epistatic genetic effects (Riedelsheimer et al., 2012). Modeling only additive effects on quantitative resistnace to stripe rust is not a major issue, nonetheless, due to previous studies showing mainly additive effects of high-temperature adult-plant (HTAP) resistance to stripe rust (Chen and Line, 1995b, 1995a). Ultimately, linear regression models assume continuous and normally distributed phenotypes, whereas machine learning models are not restricted to a certain distribution of response variable and this causes an issue on the analysis of traits (González-Camacho et al., 2018).

Data transformation is another approach used to deal with skewed and ordinal trait information. Logarithmic or square root transformations are commonly implemented to transform data for small sample sizes (Montesinos-López et al., 2015c), where they are considered standard procedures to stabilize variance, but fail to normalize inflated count data (O’Hara and Kotze, 2010; Montesinos-López et al., 2015b). Moreover, transforming data results in a loss of accuracy and power in models, especially in a small sample size (Montesinos-López et al., 2015a). When transformations are used on count data with a high number of zeros causing overdispersion, transformations may not be able to create a normal distribution (Montesinos-López et al., 2016). Another issue with using transformations is the resulting negative predicted values which are not plausible for disease resistance scores.

Another approach is to use GLMs, which accommodate non-normal data with heterogenous variance and correlated observations (Montesinos-López et al., 2015b, 2015a). GLMs provide more sensible results and have greater power to identify model effects as statistically significant (Montesinos-López et al., 2015b). Poisson and negative binomial regression models are the most common GLMs used for count and ordinal data (Montesinos-López et al., 2015c). GLMs model a function of the response mean as a linear function of the coefficients rather than modeling *y* as a linear function. These models have advantages over linear models due to their ability to model a skewed non-negative discrete distribution towards lower numbers as seen in disease resistance phenotypes (Montesinos-López et al., 2016). Several studies have shown the feasibility of integrating GLM parametric approaches into GS models such as Bayesian logistic ordinal regression (BLOR), threshold genomic best linear unbiased predictor (TGBLUP), and Bayesian mixed-negative binomial (BMNB) genomic regression (Montesinos-López et al., 2015b, 2015a, 2015c, 2016) and observed that the ordinal models present a viable alternative for predicting ordinal traits.

The last approach is to use machine learning algorithms, and classification models with no restriction on the distribution of response variables are less sensitive when modeling ordinal scores while also accounting for epistatic effects (Ornella et al., 2014; González-Camacho et al., 2018). Support vector machines (SVM) previously displayed higher performance for relative efficiency and kappa than traditional regression models such as Bayesian LASSO, Ridge Regression, and Reproducing Hilbert spaces (Ornella et al., 2014; González-Camacho et al., 2018). For the classification models, Ornella et al. (2014) further showed the superiority of SVM as the best performing model compared to random forest (RF). Additionally, classification models displayed an advantage in selecting the top performing lines.

Resistance to diseases, such as stripe rust (caused by *Puccinia striiformis* Westend. f. sp. *tritici* Erikss.) in wheat (*Triticum aestivum* L.) is commonly recorded in ordinal scales and percentages that do not follow the assumptions of linear regression models (Montesinos-López et al., 2015a; González-Camacho et al., 2018). The unbalanced, skewed distribution of resistant phenotypes is another issue for disease resistance traits in breeding programs. For example, in most wheat breeding programs, disease resistance is selected early (i.e., headrow selection before yield trials) in the breeding process. Consequently, this early selection and screening process skews the lines in disease nurseries and yield trials towards mostly resistant lines. Therefore, not only are disease-resistant traits commonly expressed in ordinal and categorical scales, but they can also be very skewed towards resistance and no longer follow a normal distribution.

Stripe rust is one of the most devastating diseases of wheat worldwide (Chen, 2020) and is especially destructive in the western United States (Chen and Line, 1995b; Rutkoski et al., 2014; González-Camacho et al., 2018; Liu et al., 2019) causing more than 90% yield losses in fields planted with susceptible cultivars (Liu et al., 2020). The use of resistant varieties and fungicide applications are the primary methods to control stripe rust (Chen and Line, 1995b; Liu et al., 2020). Quantitative stripe rust resistance, also known as adult-plant resistance (APR) or HTAP resistance, which is usually a non-race specific resistance associated with durable resistance with some genes being effective for more than 60 years (Klarquist et al., 2016).APR is conferred by different numbers of loci with varying effects and often displays partial resistance, which makes it difficult to incorporate into new cultivars (Liu et al., 2019). Therefore, APR must be improved over multiple cycles of selection and can be approached similarly to other agronomic traits (Rutkoski et al., 2014; Poland and Rutkoski, 2016; González-Camacho et al., 2018). GS approaches would be able to capture the additive effects of APR and are therefore relevant for accumulating favorable alleles for rust resistance (Rutkoski et al., 2014; Michel et al., 2017).

However, most GS studies treat disease resistance as continuous values and utilize regression models and transformations for prediction whereas only a few studies have used classification methods (Ornella et al., 2012, 2014; Rutkoski et al., 2014; Muleta et al., 2017). Therefore, this study presents empirical research to (1) evaluate GS methods using all transformations, GLMs, and non-parametric models for handling ordinal categorical phenotypes; and (2) implement these methods into selected and unselected training populations for predicting stripe rust resistance. This study identified the most accurate methods for dealing with complex phenotypes in the context of disease resistance in winter wheat.

## 2 Material and Methods

### 2.1 Phenotypic Data

The Washington State University (WSU) Winter Wheat Breeding Program takes stripe rust notes every year to select for stripe rust resistant lines. Two training populations were used to compare the different methods. The first training population consists of F_3:5_ breeding lines (BL) and doubled-haploid (DH) un-replicated trials in Pullman and Lind, WA planted in 2016-2018 and 2020 growing seasons evaluated for stripe rust responses (Table 1). Due to the un-replicated nature of the single plots, each trial in the BL consisted of unique lines which resulted in a total of 2,634 lines (1,009 in Lind and 1,625 in Pullman) over all years and locations. The BL population was subjected to stripe rust resistance screening and culling in headrows the previous year in un-replicated trials and therefore, represents our prior selected population. The second training population consisted of a diverse association mapping panel (DP) with 475 lines evaluated in un-replicated trials in Central Ferry and Pullman, WA from 2013 to 2016. The DP comprised of varieties from various breeding programs in the Pacific Northwest region of the US and represented our unselected population.

**Table 1.** Study populations for stripe rust infection type and disease severity for the diversity panel (DP) and breeding line (BL) training populations phenotyped from 2013 to 2016 and 2016-2020, respectively.

| Location | Trial <sup>a</sup> | Year | Individuals | IT <sup>b</sup> 1 | SEV <sup>c</sup> 1 | IT 2 | SEV 2 | IT 3 | SEV 3 |
| --- | --- | --- | --- | --- | --- | --- | --- | --- | --- |
| Central Ferry | DP | 2013 | 475 | X | X | X | X | X | X |
| Pullman | DP | 2013 | 475 | X | X | X | X | - | - |
| Central Ferry | DP | 2014 | 475 | X | X | - | - | - | - |
| Pullman | DP | 2014 | 475 | X | X | X | X | X | X |
| Central Ferry | DP | 2015 | 475 | X | X | X | X | X | X |
| Pullman | DP | 2015 | 475 | X | X | X | X | X | X |
| Central Ferry | DP | 2016 | 475 | X | X | X | X | X | X |
| Pullman | DH | 2016 | 136 | X | X | X | - | - | - |
| Pullman | F<sub>5</sub> | 2016 | 173 | X | X | X | - | - | - |
| Lind | F<sub>5</sub> | 2017 | 171 | X | X | X | - | - | - |
| Lind | DH | 2017 | 29 | X | X | X | - | - | - |
| Pullman | DH | 2017 | 34 | X | X | X | X | X | X |
| Pullman | F<sub>5</sub> | 2017 | 506 | X | X | X | X | - | - |
| Lind | DH | 2018 | 448 | X | X | - | - | - | - |
| Pullman | DH | 2018 | 732 | X | X | X | X | X | X |
| Pullman | F<sub>5</sub> | 2018 | 65 | X | X | X | X | X | X |
| Lind | DH | 2020 | 373 | X | X | - | - | - | - |

<sup>a</sup> Trial: DP: Diversity panel; DH: Doubled-haploid
<sup>b</sup> IT: Infection type
<sup>c</sup> SEV: Disease severity
X: Indicates measurement recorded; -: No measurement recorded

The disease traits measured were stripe rust infection type (IT) and stripe rust disease severity (SEV). The IT was based on a 0-9 scale (resistant: 0-3; intermediate: 4-6; susceptible: 7-9 (Line and Qayoum, 1992), whereas SEV was measured as the percentage of the total area of the leaf infected using a modified Cobb Scale (Peterson et al., 1948). Stripe rust data were dependent on natural infection and incidence at the time of observation. Some trials had three observations and were identified with sequential numbers. The trials with only one observation were recorded right after anthesis to measure stripe rust responses at the adult-plant stage. The reason there was only one observation was that stripe rust was not present in the field at earlier growth stages. If there were three observations, stripe rust was present in the field at earlier growth stages where the first, second, and third scores were taken soon after flag leaf emergence, after anthesis, and at early milk stage, respectively. Entries with a high infection type in the first observation, but a low infection type in the following observations may indicate that they have a HTAP resistance(Chen, 2013). However, due to the nature of APR being effective in the adult stage and that not all trials had multiple recordings, only the last observation for each trial was used to measure the stripe rust response.

### 2.2 Phenotypic Adjustments

In order to compare the regression and classification strategies, we used multiple methods of phenotypic adjustments. For the regression models, standard adjusted means were calculated considering the field design used. The ability of rrBLUP, GLM, and SVM to predict the standard and transformed (square root (SQRT), LOG, and boxcox (BC) transformed) adjusted means were then compared. For the classification models, Bayesian and SVM models were used to predict the full-scale categories for IT and SEV with the standard adjustments for field design as our control values. We then reduced both traits using multiple number of classes to determine the scenario resulting in the highest accuracy for breeding program implementation.

For the field design adjustment for controls for both the regression and classification phenotypic adjustments, a two-step adjusted means method was used, in which a linear model was implemented to adjust both IT and SEV means within and across environments. Then, a mixed linear model was used to calculate genomic estimated breeding values (GEBVs; Ward et al. 2019). Adjusted means from the stripe rust data collected in the un-replicated trials were adjusted using residuals calculated for the un-replicated genotypes in individual environments and across environments using the modified augmented complete block design model (ACBD; Federer 1956; Goldman 2019). The adjustments were made following the method implemented in Merrick and Carter (2021), as follows:

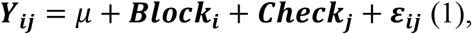

where ***Y***_***ij***_ is the phenotypic value for the trait of interest of the *i*^*th*^ block and *j*^*th*^ replicated check cultivar (i=1,…,I,j=1,…,J); *µ* is the mean effect; ***Block***_***i***_ is the fixed effect of the *i*th block; ***Check***_***j***_ is the fixed effect of the *j*^*th*^ replicated check cultivar; and ***ε***_***ij***_ are the residual errors with a random normal distribution of 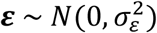. For adjusted means across environments, the model is as follows:

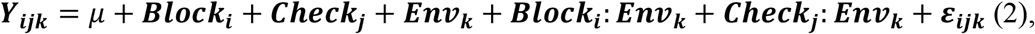

where ***Y***_***ij***_ is the phenotypic value for the trait of interest of the *i*^*th*^ block and *j*^*th*^ replicated check cultivar in the *k*^*th*^ environment (i=1,…,I, j=1,…,J, k=1,…, K); *µ* is the mean effect; ***Block***_***i***_ is the fixed effect of the *i*^*th*^ block; ***Check***_***j***_ is the fixed effect of the *j*^*th*^ replicated check cultivar; ***Env***_***k***_ is the fixed effect of the *k*^*th*^ environment; and ***ε***_***ijk***_ are the residual errors with a random normal distribution of 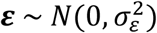.

The BLUPs for heritability were calculated for each trial and across trials using a mixed linear model for the full augmented randomized complete block design in a single environment and is as follows:

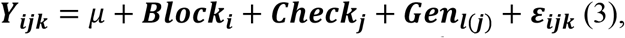

where ***Y***_***ijk***_ is the phenotypic value for the trait of interest of the *l*^*th*^ unreplicated genotype nested in the *j*^*th*^ replicated check cultivar of the *i*^*th*^ block (i=1,…,I, j=1,…,J, k=1,…,K); *µ* is the mean effect; ***Block***_***i***_ is the random effect of the *i*^*th*^ block with the distribution 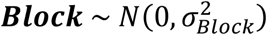; ***Check***_***j***_ is the fixed effect of the *j*^*th*^ replicated check cultivar; ***Gen***_***l*(*j*)**_ is the unreplicated genotype *l* in the *j*^*th*^ check with the distribution 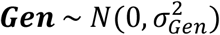; and ***ε***_***ij***_ are the residual errors with a random normal distribution of 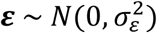. The full model across environments is as follows:

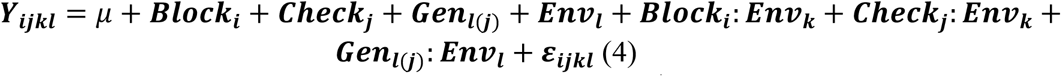

where ***Y***_***ijkl***_ is the phenotypic value for the trait of interest of the *l*^th^ unreplicated genotype nested in the *j*^*th*^ replicated check cultivar of the *i*^*th*^ block and in the *k*^*th*^ environment (i=1,…,I, j=1,…,J,k=1,…,K, l=1,…,L); *µ* is the mean effect; ***Block***_***i***_ is the random effect of the *i*^*th*^ block with the distribution 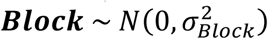; *Check*_*j*_ is the fixed effect of the *j*^*th*^ replicated check cultivar; ***Gen***_***l*(*j*)**_ is the random effect of the genotype *l* in the *j*^*th*^ replicated check cultivar with the distribution 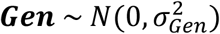; ***Env***_***k***_ is the random effect of the *k*^*th*^ environment with the distribution 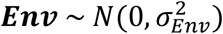; and ***ε***_***ijkl***_ are the residual errors with a random normal distribution of 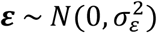. After adjustments were made, values outside of the 0-10 (IT) and 0-100 (SEV) scales were rounded back to 0-10 and 0-100, respectively, to avoid negative values for log transformations or Poisson distributions and to have the standard adjusted means for all comparisons.

Broad-sense heritability on a genotype-difference basis was calculated using the variance components from the models (3) and (4) implemented Merrick and Carter (2021) and using BLUP for both individual environments and across environments (Cullis et al., 2006):

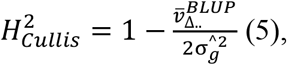

where 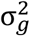, and 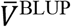 is the genotype variance and mean-variance of a difference between two BLUPs for the genotypic effect BLUPs, respectively (Schmidt et al., 2019). Trial evaluations were compared using general summary statistics, coefficient of variations (CV), skewness, kurtosis and the non-parametric Kruskal-Wallis test using the R package “ggpubr” (R Core Team, 2018; Kassambara and Kassambara, 2020).

### 2.3 Genotypic Data

Wheat lines were genotyped using genotyping-by-sequencing (GBS; Elshire et al. 2011) through the North Carolina State University (NCSU) Genomics Sciences Laboratory in Raleigh, North Carolina (https://research.ncsu.edu/gsl/) using a two-enzyme (*PstI/MspI*) digestion protocol (Poland and Rife, 2012). Genomic DNA was isolated from individual seedlings at the one-to three-leaf stage using Qiagen BioSprint 96 Plant kits and the Qiagen BioSprint 96 workstation (Qiagen, MD, USA). Genotyping by sequencing was conducted using Illumina HiSeq^®^ 2500 and NovaSeq 6000. Sequences were aligned to the Chinese Spring International Wheat Genome Sequencing Consortium (IWGSC) RefSeq v1.0 (Appels et al., 2018) using the Burrows-Wheeler Aligner (BWA) 0.7.17 (Li and Durbin, 2009). GBS derived single-nucleotide polymorphism (SNP) markers were called using TASSEL-GBS v2 SNP calling pipeline in TASSEL v5.2.35 (Bradbury et al., 2007; Glaubitz et al., 2014). Markers with > 20% missing data, minor allele frequency (MAF) <5%, and those that were monomorphic were removed. Imputation of missing genotypes were conducted using Beagle 5.0 (Browning et al., 2018) and markers with < 5% MAF were further excluded. The remaining markers were binned together based on a linkage disequilibrium threshold value of 0.80 (Ward et al., 2019). The reduced genotype matrix was computed using JMP genomics version 9 (SAS Institute, Inc, 2011). Principal components analysis (PCA) using the SNP data was performed using ‘prcomp’ and a biplot with *K*-mean clusters was created using the “autoplot” packages in R. Cluster number for *k*-means were calculated according to the elbow method using a scree plot with the optimal number of clusters identified when the total intra-cluster variation was minimized.

### 2.4 Regression Models

#### 2.4.1 Transformations

Transformations using SQRT, LOG, and BC approaches were compared to determine the optimal method for phenotypic adjustment for skewed phenotypes (Table 2). The BC transformations were conducted using the “forecast” package (Hyndman and Khandakar, 2008) that identify optimal lambda values using the “BoxCox.lambda” function in R.

**Table 2.**
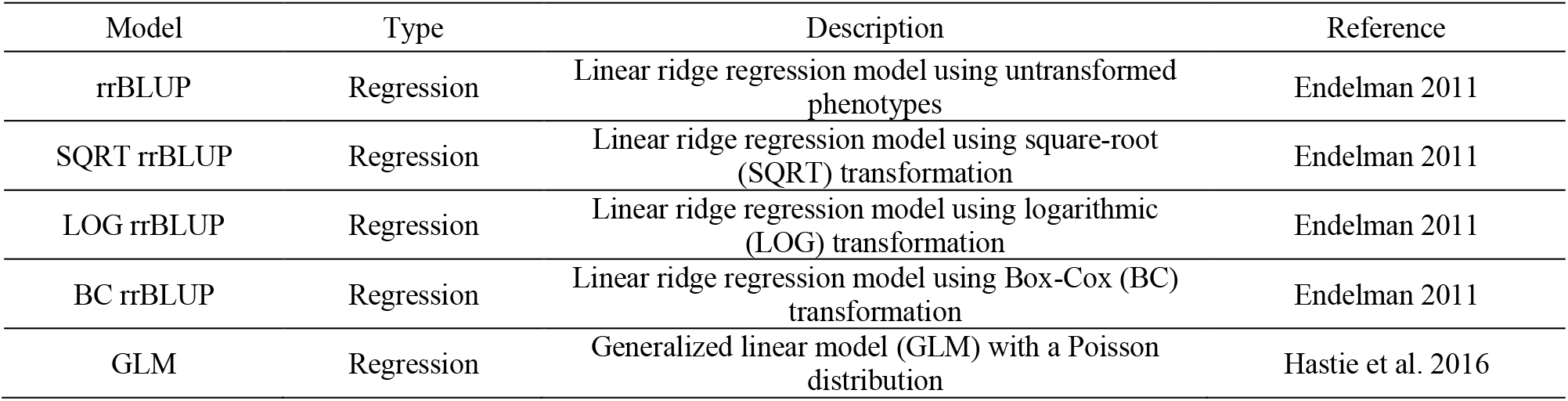
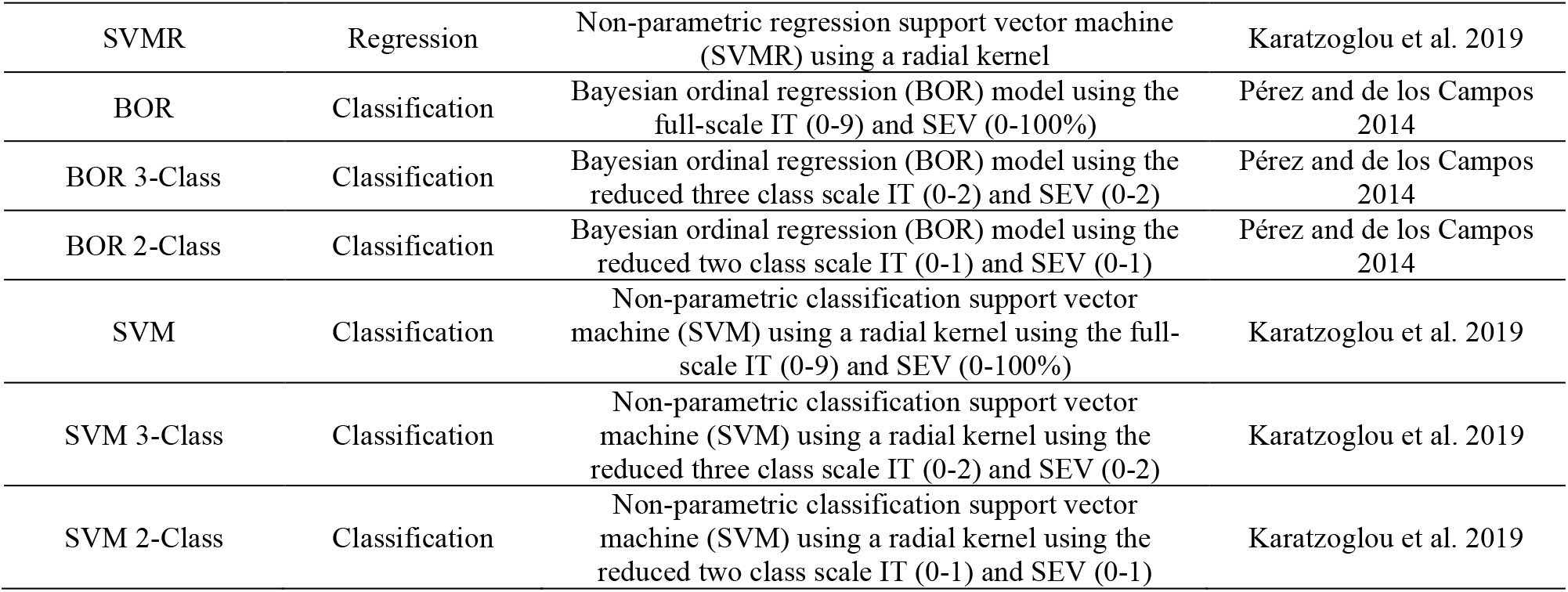
Regression and classification genomic selection models for stripe rust infection type (IT) and disease severity (SEV) in winter wheat.

#### 2.4.2 rrBLUP model

Ridge regression best linear unbiased prediction (rrBLUP) was used as the standard GS model for comparing the predictive ability of the adjusted means and transformed data. The rrBLUP was selected due to its high predictive performance for stripe rust resistance (Table 2; Rutkoski et al. 2014; Arruda et al. 2016; Poland and Rutkoski 2016; Muleta et al. 2017; Merrick et al. 2021). The model follows the basic mixed linear model that treats the effects of markers as random effects as described by Endelman (2011):

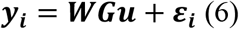

where 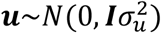is a vector of marker effects; ***y***_***i***_ is a vector of phenotypes; G is the genotype matrix; W is the design matrix for **y**. The marker effects are then calculated using ***û*** = (***Z***′***Z*** + *λ****I***)^−1^***Z***′***y*** with the ridge parameter of 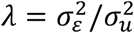, which is the ratio of the residual and marker variances.

#### 2.4.3 Generalized Linear Model

The generalized linear model (GLM) was implemented using “Glmnet” with a Poisson distribution (Table 2; Hastie et al. 2016). Glmnet fits a GLM via penalized maximum likelihood with the elastic net penalty computed at grid values on the log scale for the regularization parameter lambda. Glmnet solves the equation:

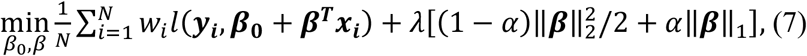

over a grid of values of *λ*; *l*(*y*_*i*_ − *η*_*i*_)^2^ is the negative log-likelihood of *i*. The elastic net penalty is controlled by *α* and bridges the game between lasso regression (*α*=1) and ridge regression (*α* = 0), with *λ* controlling the penalty. ***y***_***i***_ is a vector of phenotypes; ***β*** is the genotype matrix; ***x*** is the design matrix for **y**. Poisson regression is used to model count data under the assumption of Poisson error, or otherwise non-negative data where the mean and variance are proportional. Like the Gaussian and binomial models, the Poisson distribution is a member of the exponential family of distributions. We model its positive mean on the log scale: log *µ*(*x*) = ***β***_**0**_ **+ *β* ′ *x***.

### 2.5 Classification Models

#### 2.5.1 Factor Adjustments

We used a Bayesian ordinal model and a SVM to compare factor adjustments (Table 2). The adjusted means were used as control for categorical factors but rounded to discrete values, so they follow the initial ordinal scales for both IT and SEV. These scales are 0-9 for IT and 0-100 for SEV. The original 0-9 IT scale and 0-100 SEV scale were reduced to a three-class 0-2 scale (resistant/intermediate/susceptible), and a binary keep/discard scale of 0-1 in order to be more applicable to breeding programs and reduce the effect of unbalanced classes.

#### 2.5.2 Bayesian Ordinal Regression model

The Bayesian Ordinal Regression (BOR) model implemented in the BGLR package according to Pérez and de los Campos (2014) follows:

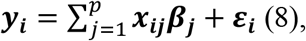

where ***y***_***i***_ is a vector of phenotypes; ***x***_***ik***_ is the genotype of the *k*^*th*^ marker and *i*^*th*^ individual, *p* is the total number of markers, ***β***_***k***_ is the estimated random marker effect of the *k*^*th*^ marker;and ***ε***_***i***_ is a vector of residuals with a random normal distribution of 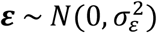. Each version of the BOR model has its own conditional prior distribution and a scaled-inverse chi-squared density described in Pérez and de los Campos (2014) whose hyper-parameters are set internally by the software. The BOR model uses the probit link function in which the probability of each of the categories is linked to the linear predictor according to the link function outlined in Pérez and de los Campos (2014):

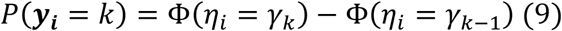

where Φ(· ) is the standard normal cumulative distribution function, *η*_*i*_ is the linear predictor, and *γ*_*k*_ are threshold parameters, with *γ*_0_ = −∞, *γ*_*k*_ ≥ *γ*_*k*−1,_ *γ*_*k*_ = ∞. The BOR model was implemented in the “BGLR” package in R with a burn-in rate of 10,000 and 80,000 iterations based on convergence of the models using trace plots (Pérez and de los Campos, 2014; Merrick and Carter, 2021).

#### 2.5.3 Support Vector Machine

The Support Vector Machine (SVM) is a non-parametric model that can be used for both classification and regression (SVMR) with no specific phenotypic distribution requirement. The SVM performs well in a variety of settings due its use of a maximal margin classifier. The maximal margin classifier uses a hyperplane to classify and separate observations by computing the maximum distance of an observation to the hyperplane and then determining the class of the observation based on which side of the hyperplane it falls on (Gareth et al., 2013). Additionally, SVMs can enlarge the feature space of the data using kernels to accommodate non-linear boundaries between classes and simplifies the inner product, which overcomes the dimensionality of the data. For classification, the radial basis function (RBF) was used due to its wide adaption and ability to be applied to any distribution of observations (Wang et al., 2018). Both SVM and SVMR were implemented using the “caret” package in R, with the RBF model using the “kernlab” function in R (Kuhn, 2008; Karatzoglou et al., 2019; Meyer et al., 2019). Further, model tuning was completed using five replications of ten-fold CV with resampling within the training set of the training fold of the cross-validation or validation sets. Additionally, for classification, the SVM model was tuned using up-sampling which randomly samples the minority class to be the same size of the majority class in order to deal with class imbalances which can have significant negative impact on model fitting (Kuhn, 2008).

### 2.6 Prediction Accuracy and Scheme

Prediction accuracy for the regression models was reported using Pearson correlation coefficients (*r*) and prediction bias was reported using root mean square error (RMSE) between GEBVs and their respective adjusted means using the function “cor” in R. However, due to the unbalanced class type, the classification models were evaluated using overall class accuracy (*R*^*2*^) using the “confusionMatrix” function in the “caret” package and reported as *R*^*2*^ (Kuhn, 2008). Cohen’s kappa coefficient (kappa) was used to evaluate classification model bias because it takes into account unbalanced classes (Ornella et al., 2014; González-Camacho et al., 2018).

In order to compare regression and classification models, relative efficiency (RE) was used. RE is based on expected genetic gain when individuals are selected by GS compared to the individuals selected by phenotypic selection. The model for RE according to Ornella et al. (2014) is:

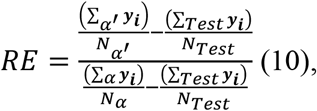

where *α* and *α*′ are the to 15% of individuals selected by the ranking of observed or predicted values, respectively. *N*_*α*_ = *N*_*α*_′ is the number of individuals selected; ***y***_***i***_ is the observed phenotypic value of the *i*th individual; and 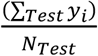 is the mean of the test population. The denominator is the selection differential of the individuals selected by phenotypic selection and the numerator is the selection differential of the individuals selected by GS. The 15% selection intensity was chosen due to its performance of RE when replacing phenotypic selection with GS (Ornella et al., 2014; González-Camacho et al., 2018).

The prediction accuracy was assessed using a five-fold cross-validation scheme and independent validation sets for IT and SEV in the DP and BL training populations (Merrick et al., 2021). The two populations were used to compare the effects of a selected and unselected population with varying degrees of resistance. Models for GS were conducted with five-fold cross-validation by including 80% of the samples in the training population and predicting the GEBVs of the remaining 20% (Merrick and Carter, 2021; Merrick et al., 2021). One replicate consisted of five model iterations, where the population was split into five different groups.

Independent validation sets were then performed according to Merrick and Carter (2021) on a yearly basis by combining the two locations for each training population and predicting the following year which results in three continuous training scenarios for each population. For example, the combination of Pullman and Central Ferry trials for the DP in 2013 were used as a training population to predict the combination of Pullman and Central Ferry trials in the DP in 2014. Final validation set was completed by combining all years and locations within a training population and then predicting the combination of years and locations for the other training population. All trials in the BL in both Pullman and Lind combined across 2016 to 2020 was used to predict all trials in the DP in both Central Ferry and Pullman across 2013 to 2016. This allows the evaluation of models in a realistic breeding situation in which we combine all available data to build a training population. All cross-validations and independent validations were replicated 10 times. All GS and MAS models and scenarios were analyzed using WSU’s Kamiak high performance computing cluster (Kamiak, 2021). Model, scenario, and training population comparisons were evaluated by using a Tukey’s honestly significant difference (HSD) test implemented in the “agricolae” package in R (de Mendiburu and de Mendiburu, 2019). The comparison of models was then plotted for visual comparison using “ggplot2” in R (Wickham, 2011).

## 3 Results

### 3.1 Phenotypic Data

The stripe rust phenotypes for both IT and SEV demonstrated variability for each scale (Table 3). For the DP, the IT and SEV values ranged the entire scale of each trait for the majority of the trials. Additionally, the means of the DP were higher than the BL trials, with lower coefficient of variations (CV). Further, the BL trials ranged the entire scale for IT, but had lower means. The SEV in the BL trials did not reach the maximum value of SEV. Overall, the BL displayed a higher proportion of resistance than the DP trials. Every trial and trait displayed a positively skewed distribution, with the exception of SEV in the DP in Pullman in 2015. SEV for the majority of trials were extremely skewed for the BL, with Lind in 2018 displaying the highest skew of any trial and trait. Skewness decreased for combined analysis across environments. Positive values above 3 display long skinny tails as in the case for SEV for the BL population in Lind (2018) at 19.77. The majority of distributions are skinny tailed demonstrating the large amount of similar disease resistance around 0 and the large amount of resistance in the BL and DP populations.

**Table 3.**
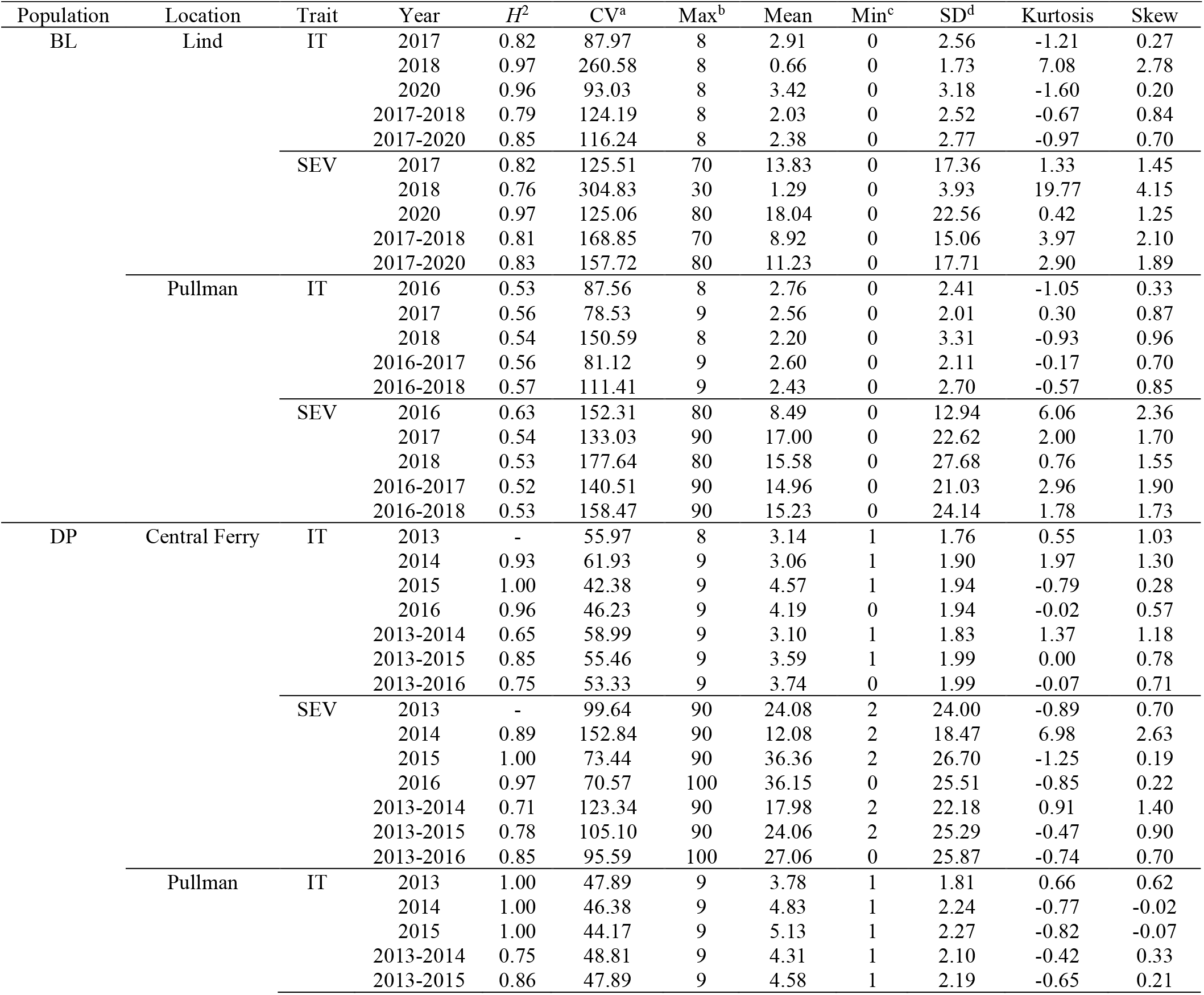
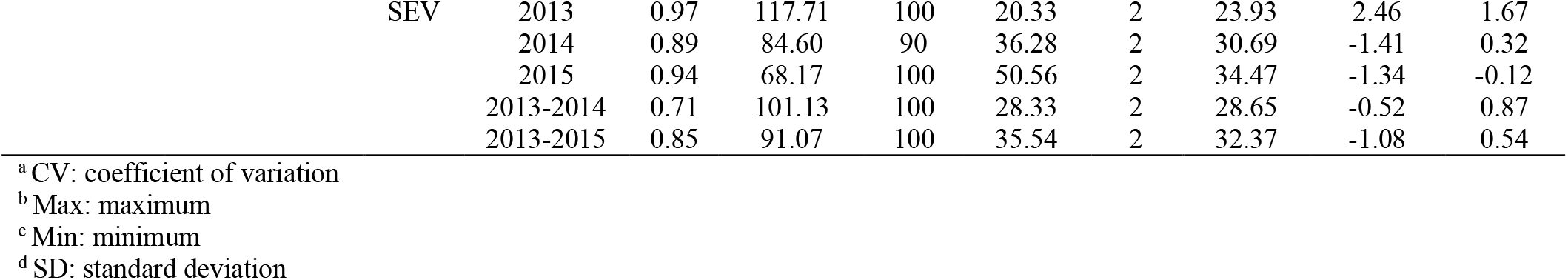
Stripe rust infection type (IT) and disease severity (SEV) heritability (*H*^*2*^) and trial statistics for unadjusted phenotypes in the diversity panel (DP and breeding line (BL) training population phenotypes from 2013 to 2016 and 2016 to 2020 growing seasons.

The skewness and kurtosis of the distributions were further visualized (Figure 1). The DP is less skewed than the BL. For both IT and SEV, the DP displayed more variation than the BL, except for SEV in Central Ferry. Further, there were significant differences between most years for each population and location (Figure 1). Heritability of the BL trials were moderately high for both IT and SEV, with values ranging from 0.76-0.97 and 0.52-0.63, respectively. For the DP, heritability ranged from 0.65-1.00 for IT and 0.71-1.00 for SEV (Table 1).

**Figure 1.**
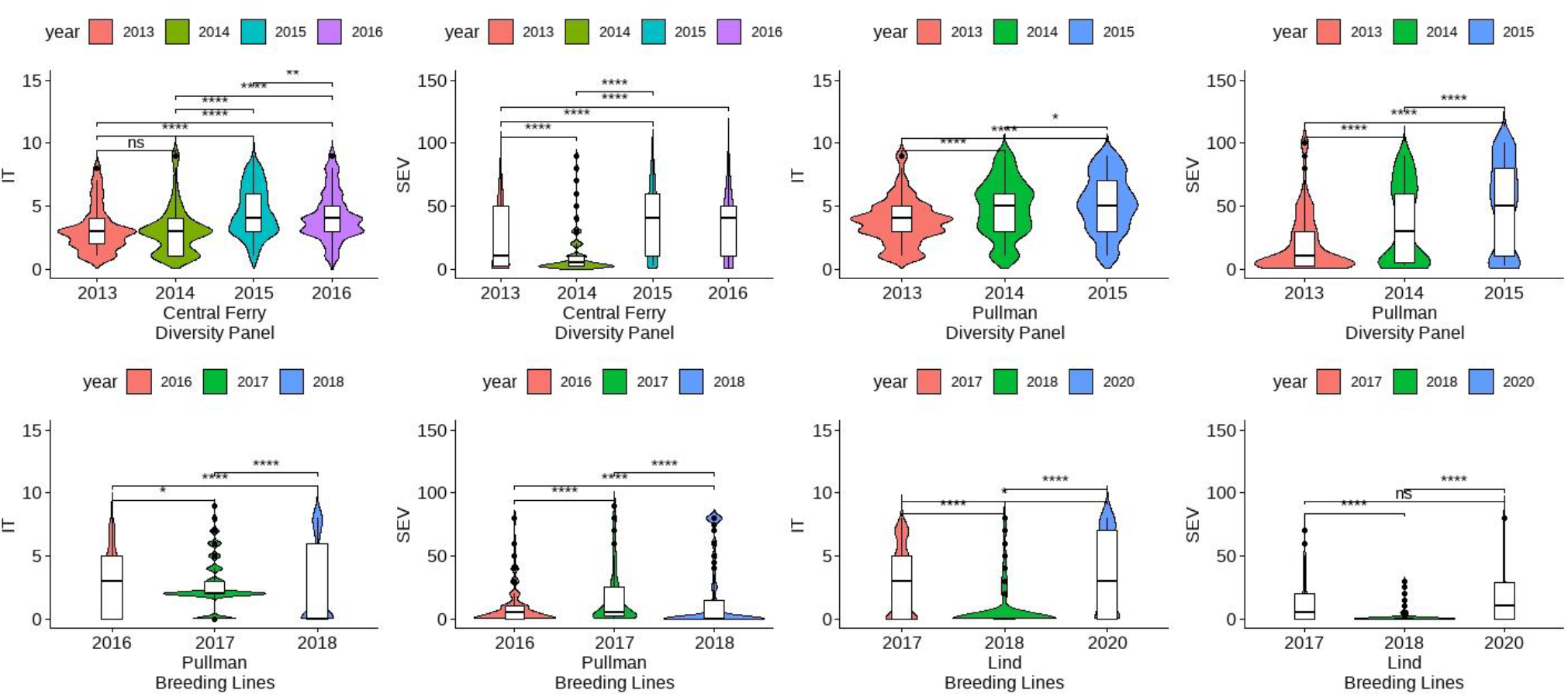
Comparison of infection type (IT) and disease severity (SEV) over years and locations in the diversity panel and breeding line training populations using Kruskal-Wallis test. Models labeled with the same letter are not significantly different.

### 3.2 Analysis of principal components

After filtering and imputation, a total of 19,861 SNP markers for the 475 unique DP lines and the 2,630 BL lines were obtained from GBS. Principal component analysis using SNP markers for the DP and BL populations resulted in four clusters with Cluster 2 (green) overlapping with the other clusters (Figure 2). PC1 explained 5.8% of the variation whereas PC2 explained 3.4% of the variation. The biplot displayed four main clusters over the combined populations using *k*-means clustering. Cluster 1 comprised of lines common in both the BL and DP. Majority of lines in both the DP and BL were included in Cluster 3, which comprised of BL in Lind and Pullman and lines from the DP. Cluster 4 consisted mainly of lines from the BL in Lind, whereas majority of lines from the BL in Pullman comprised Cluster 2.

**Figure 2.**
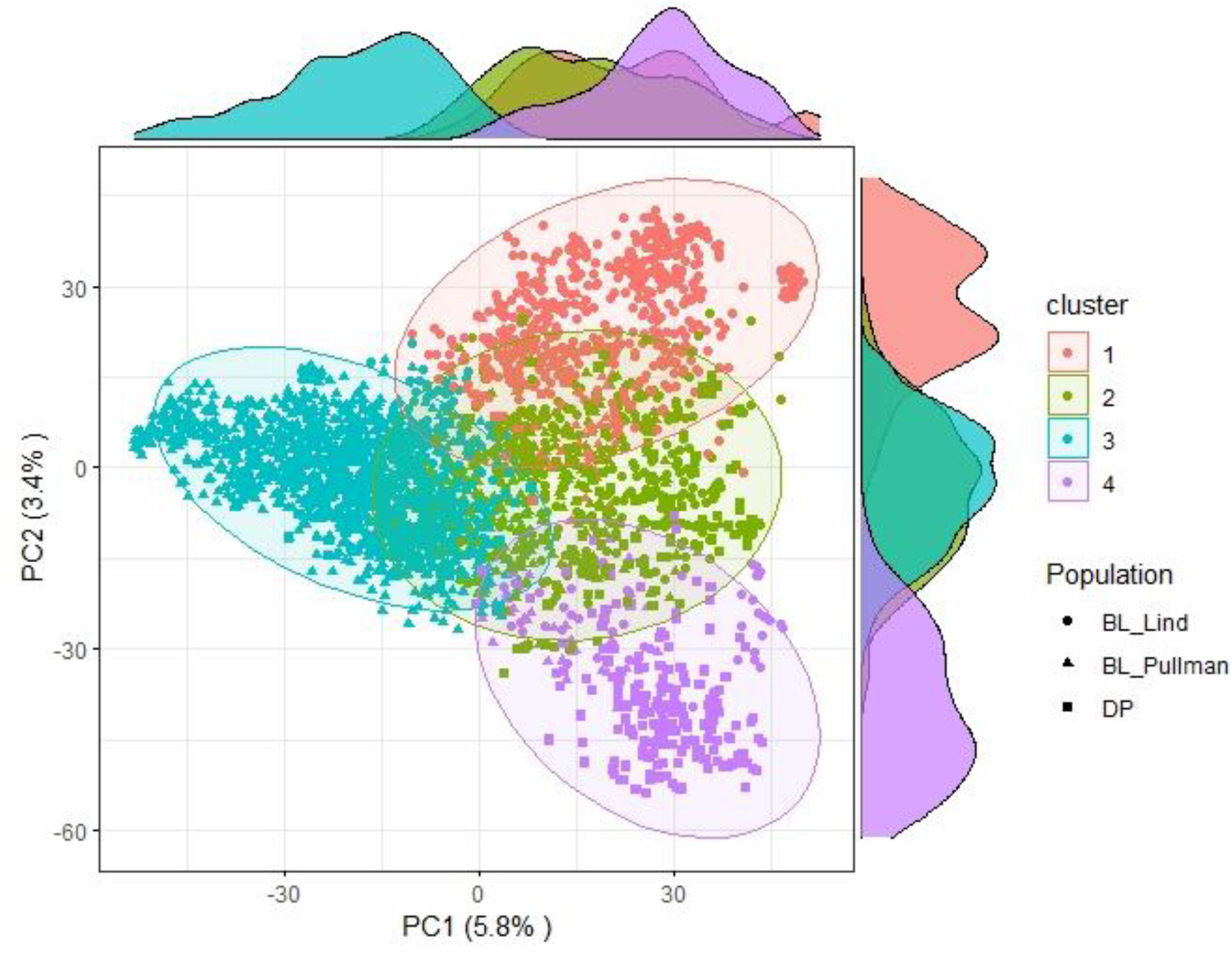
Principal component (PC) biplot and *k*-means clustering of SNP GBS markers from the diversity panel and breeding line training populations.

### 3.3 Cross-validations for regression models

Multiple comparisons for RMSE and Pearson correlations for accuracy were conducted for the regression models in individual populations and years for IT and SEV. The SVMR model resulted in the highest accuracy (*r*= 0.73) in the 2018 Pullman BL trial for IT (Figure 3). Accuracy for the GLM model in 2018 Pullman BL was 0.72. The GLM displayed consistent high accuracies in the more skewed BL population than the less skewed DP but displayed the lowest accuracy for the most skewed trial in the BL in Lind in 2018 (0.23). Overall, there were no significant differences for the BL, whereas the LOG rrBLUP and the GLM model showed significant differences (*P* < 0.05) in the DP. Additionally, the BL trials had higher mean accuracies than the DP trials with an increase in accuracy with the combination of years. Altogether, the rrBLUP had the highest accuracy over the transformed phenotypes (0.53). The rrBLUP model had similar RMSE than the SVMR and GLM models with 2.15, 2.18, and 2.28, respectively (Figure S1). The SQRT rrBLUP model had the lowest RMSE (0.51), and the BC and LOG rrBLUP models had the highest RMSE (5.67 and 5.93, respectively). Using SQRT transformation on the phenotypes reduced the error of the predictions compared to the other transformations.

**Figure 3.**
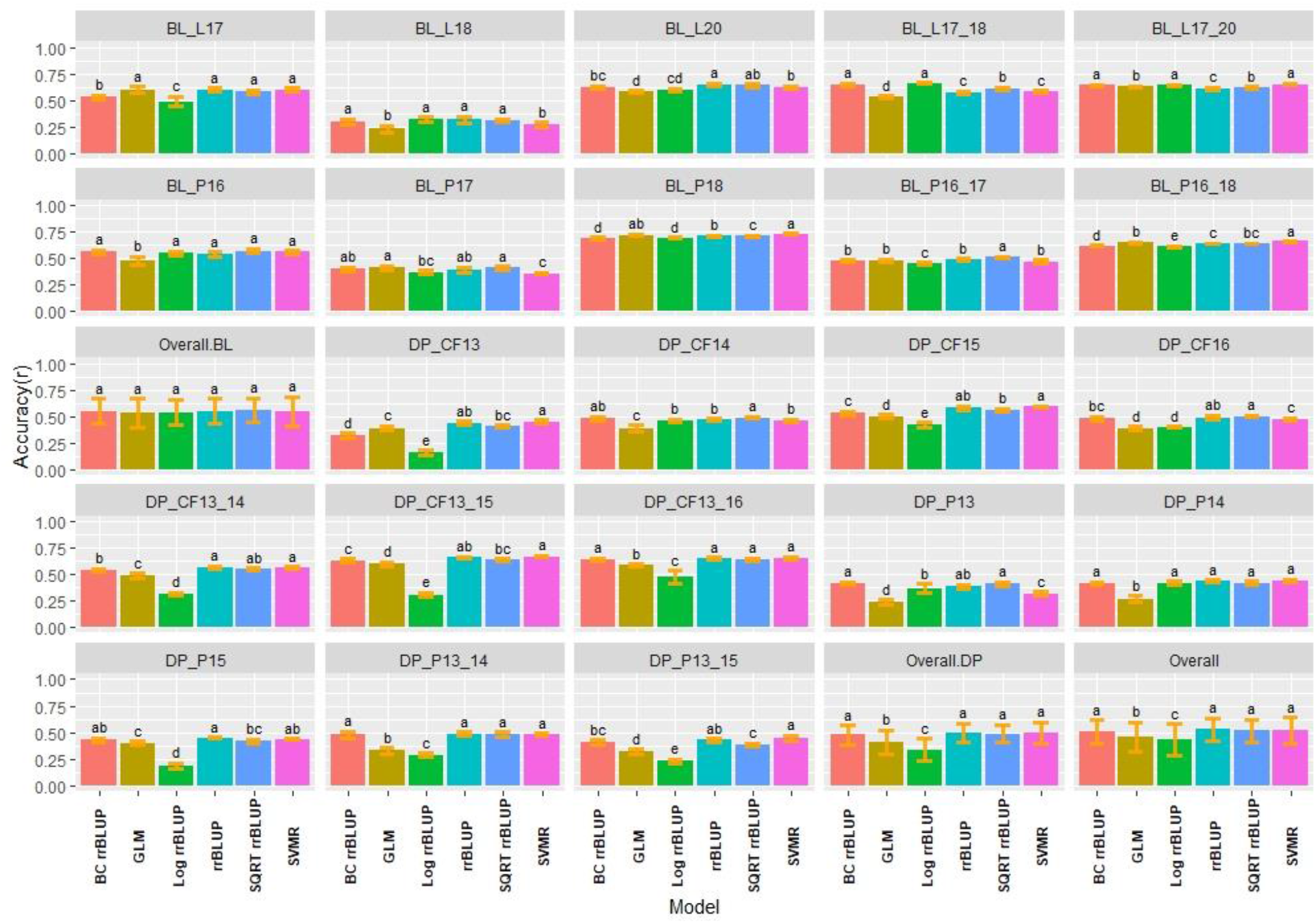
Pairwise comparisons of genomic selection regression model accuracy (*r*) using cross-validations for stripe rust infection type. Pacific Northwest winter wheat diversity panel (DP) lines phenotyped from 2013 to 2016 in Central Ferry (DP_CF) and Pullman (DP_P), WA. Washington State University breeding lines phenotyped from 2016 to 2020 in Lind (BL_L) and Pullman (BL_P), WA. Model comparison across the DP (Overall.DP), BL (Overall.BL), and Overall scenarios. Models labeled with the same letter are not significantly different. Bars indicate standard errors.

Similar to IT, the highest accuracies for SEV were obtained in the 2018 Pullman BL trial, with the GLM reaching the highest accuracy (0.76), followed by the SQRT rrBLUP (0.74) and SVMR (0.73) models (Figure S2). The lowest accuracies were also achieved with the GLM model in the 2018 Lind BL trial (0.18). The 2018 Lind BL trial had the lowest accuracies for the majority of models. Similar to IT, there were no statistical differences between models overall in the BL, and the SQRT rrBLUP, rrBLUP, and SVMR reached the highest accuracies in the DP. The SVMR and SQRT rrBLUP reached the highest accuracies of 0.60 (Figure S2). For SEV, the RMSE for the transformed rrBLUP models displayed much lower RMSE values than the rrBLUP, GLM, and SVMR models (Figure S3). However, this discrepancy is presumably due to the phenotypic range of the transformations compared to the untransformed range for SEV, which is 0-100. The BC rrBLUP model displayed an extremely large RMSE in the DP in Central Ferry in 2015 (57.11). Overall, the rrBLUP models displayed statistically similar RMSE values with the transformed rrBLUP models.

### 3.4 Cross-validations for classification models

Due to the difference between regression and classification models, multiple comparisons for the kappa coefficient and overall class accuracy were conducted for the classification models in individual populations and years for IT and SEV. In contrast to the regression models where the 2018 Lind BL trial had the lowest regression accuracies, the classification models displayed the highest *R*^*2*^ values with the 2-class and 3-class BOR models reaching an overall class accuracy of 0.88 for IT (Figure 4). Additionally, the SVM models displayed much higher accuracies than the BOR models overall. The full scale BOR model had very low accuracy for the majority of trials with the BL in 2018 in Pullman. The reduced class sizes, 2 and 3, displayed higher accuracy than the full IT scales. Overall, the selected BL displayed higher accuracies than the unselected DP. The 2-Class SVM reached the highest overall class accuracy with 0.76 in the BL and 0.69 in the DP. The 2-Class SVM reached the highest overall class accuracy of 0.72 in the overall comparison. The high-class accuracies in the BL in Lind in 2018 can be explained by the kappa values of 0 (Figure S4), displaying the highly skewed data and the inability for the models to account for phenotypes of mostly zeros. The SVM displayed lower kappa values in the DP than in the BL, but the BOR models had the opposite trend. The BOR models displayed higher kappa values than the SVM models, but the SVM models showed higher accuracy.

**Figure 4.**
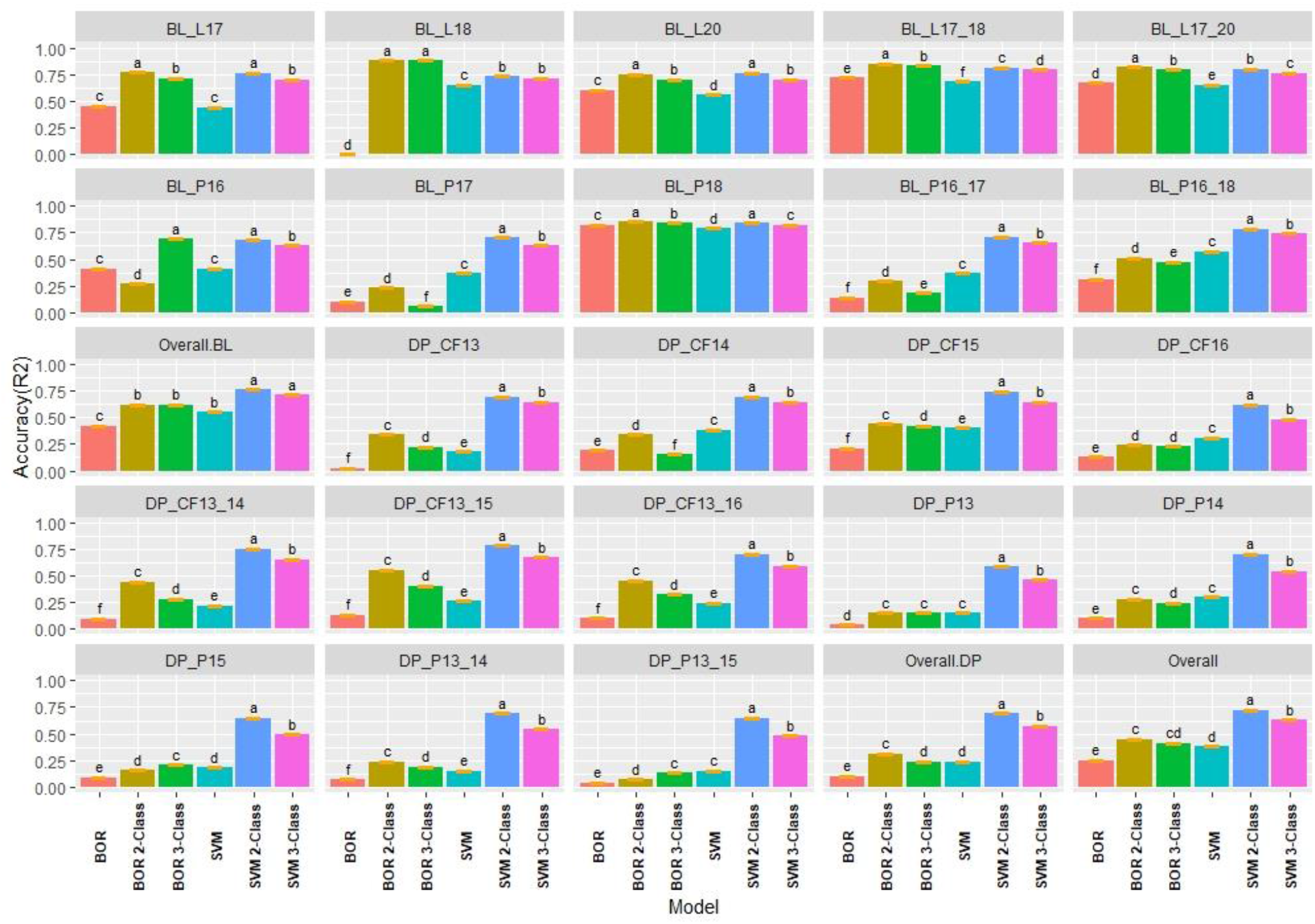
Pairwise comparisons of genomic selection classification model overall class accuracy (*R*^*2*^) using cross-validations for stripe rust infection type. Pacific Northwest winter wheat diversity panel (DP) lines phenotyped from 2013 to 2016 in Central Ferry (DP_CF) and Pullman (DP_P), WA. Washington State University breeding lines phenotyped from 2016 to 2020 in Lind (BL_L) and Pullman (BL_P), WA. Model comparison across the DP (Overall.DP), BL (Overall.BL), and Overall scenarios. Models labeled with the same letter are not significantly different. Bars indicate standard errors.

The classification models for SEV had very similar results to IT, with the BOR and SVM 2-class models reaching an accuracy of 0.99 and 0.98, respectively (Figure S5). This was due to the very skewed and high levels of zeros in the data in the BL in Lind in 2018. Additionally, in the DP which had less skewed phenotypes, the BOR models showed very poor overall class accuracy with the majority of trials having R^2^ values of 0.20, with moderate accuracies for the 2-Class BOR. The 2-Class SVM displayed the highest statistically significant class accuracy in scenarios with *R*^*2*^ values of 0.86, 0.78, and 0.81 within the BL, DP, and overall comparisons, respectively. The kappa values were higher in the DP trials due to less skewed phenotypes and displayed low values in the high accuracy trial of the BL in 2018 Lind. Overall, the 2-class SVM had the highest kappa value for SEV with 0.46 (Figure S6).

### 3.5 Cross-validation relative efficiency

Relatively efficiency (RE) was used to compare the selection differential between the GS models and phenotypic selection for the phenotypes. Overall, the highest relative efficiencies for IT were the regression models with the majority of models having statistically similar relative efficiencies. The regression models had very high RE values with the rrBLUP models reaching a maximum value of 0.94 in the 2018 Pullman BL trial (Figure 5). The SVMR model had statistically similar RE values to the rrrBLUP models in the overall comparisons. In contrast, the classification models had relatively low RE in the majority of trials with the 3-class BOR model (−0.38) in the combined 2017 to 2018 Lind BL trials. This confirmed the bias seen in the kappa results with the majority of lines being predicted as zeros. Interestingly, the 2 and 3-class BOR and SVM displayed lower RE values overall than the full-scale models. Overall, the rrBLUP and SQRT rrBLUP reached RE values of 0.62.

**Figure 5.**
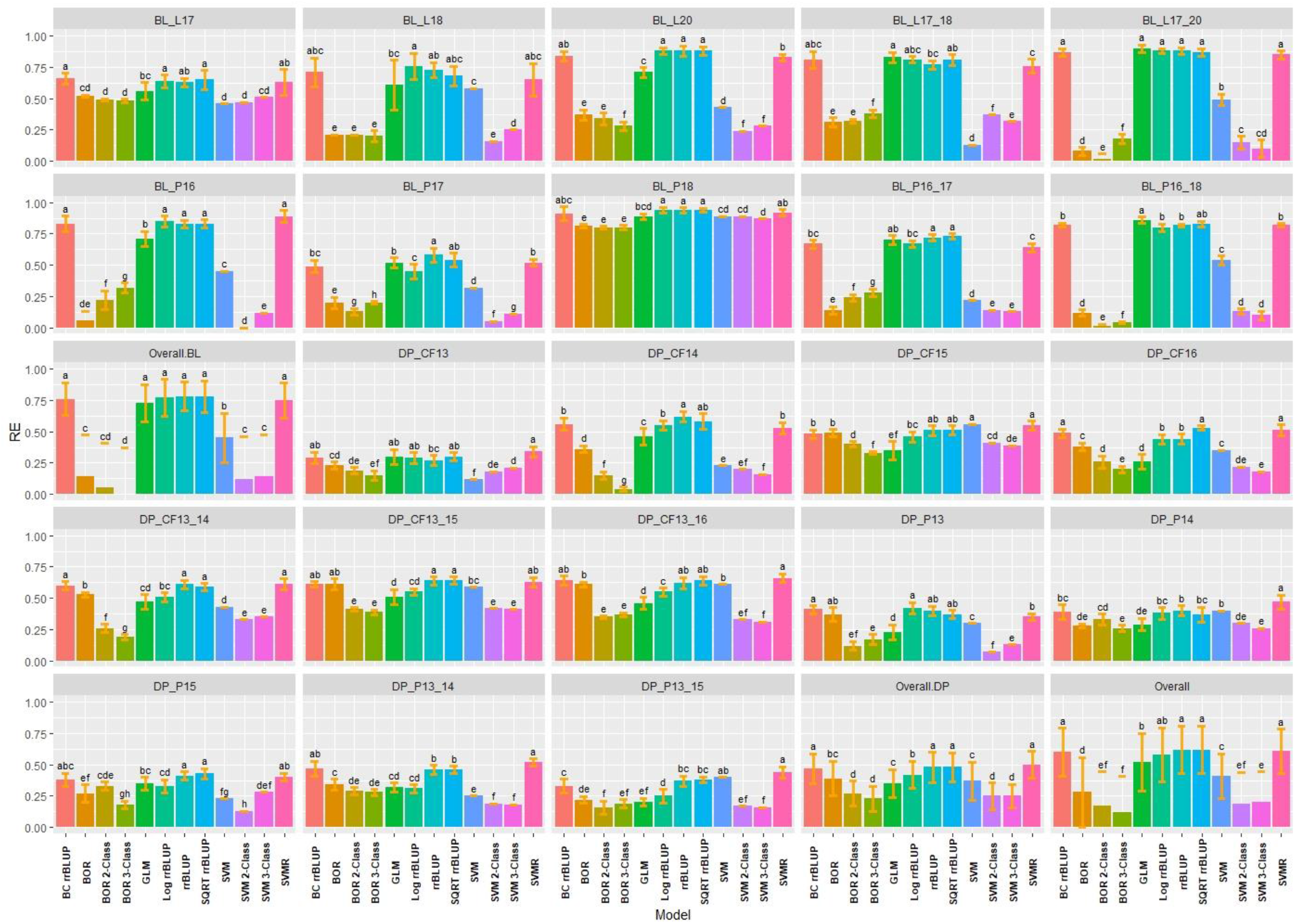
Pairwise comparisons of genomic selection regression and classification model relative efficiency (RE) using cross-validations for stripe rust infection type. Pacific Northwest winter wheat diversity panel (DP) lines phenotyped from 2013 to 2016 in Central Ferry (DP_CF) and Pullman (DP_P), WA. Washington State University breeding lines phenotyped from 2016 to 2020 in Lind (BL_L) and Pullman (BL_P), WA. Model comparison across the DP (Overall.DP), BL (Overall.BL), and Overall scenarios. Models labeled with the same letter are not significantly different. Bars indicate standard errors.

Similar to IT, the regression models had very high RE for SEV, with the classification models reaching low to moderate values ranging between −0.58 to 0.89 (Figure S7). The rrBLUP models had very high RE (0.98; BL Pullman 2018) compared to phenotypic selection. The rrBLUP models showed consistently higher REs than the GLM and SVMR models. The GLM displayed similar RE values in the BL, but lower in the DP. The SQRT rrBLUP model had the highest RE overall (0.81). The classification models had very low RE except the BL trials in 2018 Pullman and 2017 Lind, which showed very high RE compared to the other years and populations. Additionally, the combined trials for both the BL and DP displayed higher RE than some of the individual years indicating an advantage of combining trials.

### 3.6 Validation sets for regression models

The training populations were evaluated for validation sets on a yearly basis and over combined years and trials. We used the earliest trial to predict the following year and then a new model with the addition of each subsequent trial to evaluate genotype-by-environment interaction of a prediction model. We then compared the combination of all trials for one population to predict the combination of all trials in the other population. The highest accuracy for IT was in the continuous training scenario of the DP combined 2013-2015 to predict the DP 2016 with SQRT rrBLUP reaching 0.65 (Figure 6). There were only a few significant differences, with none in the overall BL or DP. Overall, the SQRT rrBLUP displayed the highest accuracy (0.46). Further, there was an increase in accuracy as the years were combined within the same population. However, the accuracy was much lower when predicting into the combined trials of the other population. Similar RMSE values to the cross-validations were displayed with SQRT rrBLUP having the lowest RMSE (1.31; Figure S8).

**Figure 6.**
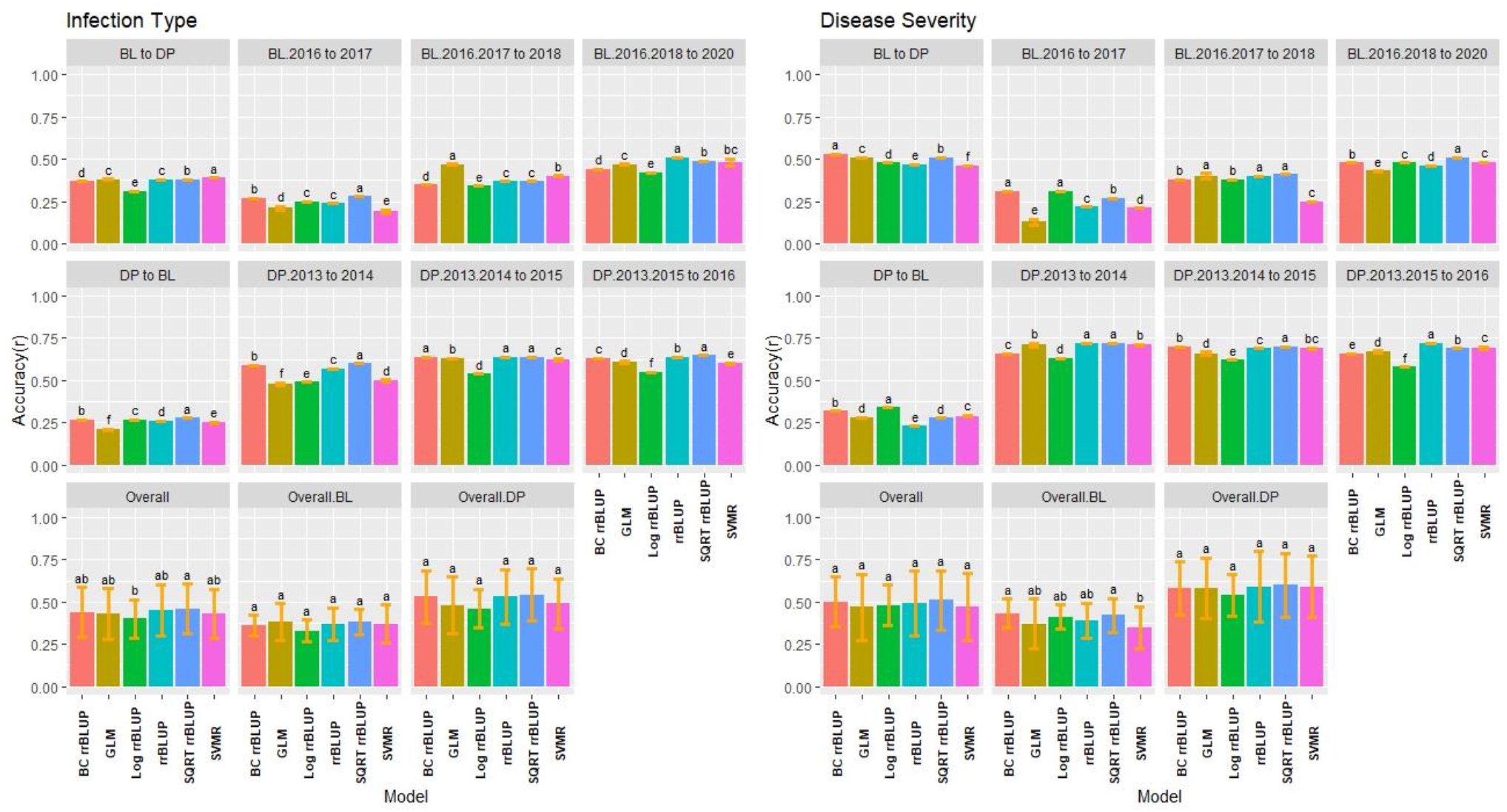
Pairwise comparisons of genomic selection regression model accuracy (*r*) using validation sets for stripe rust infection type and disease severity. Pacific Northwest winter wheat diversity panel (DP) lines phenotyped from 2013 to 2016 in Central Ferry (DP_CF) and Pullman (DP_P), WA. Washington State University breeding lines phenotyped from 2016 to 2020 in Lind (BL_L) and Pullman (BL_P), WA. Model comparison across the DP (Overall.DP), BL (Overall.BL), and Overall scenarios. Models labeled with the same letter are not significantly different. Bars indicate standard errors.

The validation accuracy for SEV displayed similar trends to IT, with the highest accuracy of 0.72 for the SQRT rrBLUP and rrBLUP. (Figure 6). Interestingly, the combined BL trials predicting into the combined DP displayed the highest accuracy in the BL prediction scenarios with BC rrBLUP reaching 0.53. This trend was in contrast to IT. However, the opposite was seen in the DP. The validation set accuracy for the DP was higher than the validation sets for BL. In the overall comparison, there were no statistical differences between the models. The BC and Log rrBLUP displayed the highest accuracies in some scenarios, which was not seen in cross-validations and was only observed in the BL. The RMSE values were much higher for SEV with the SQRT rrBLUP displaying similar RMSE to IT (Figure S9).

### 3.7 Validation sets for classification models

The classification models had contrasting results for the validation sets compared to the regression models. The validation set class accuracy for the classification models were all relatively low except the two and three class SVM model. Furthermore, there was no trend in increasing overall class accuracy by combining trials. The BL trials displayed the highest overall class accuracy with *R*^*2*^ values reaching 0.78 for the two class SVM model (Figure 7). The low accuracies were presumably due to the increase in resistance and the models predicting zeros in the IT scale. Similar to the cross-validation scenarios, the reduced two class models reached a much higher accuracy across the majority of trials. Further, the prediction accuracy can be accounted for by the low kappa values in the majority of models except the 2-Class SVM model reaching 0.40 (Figure S10).

**Figure 7.**
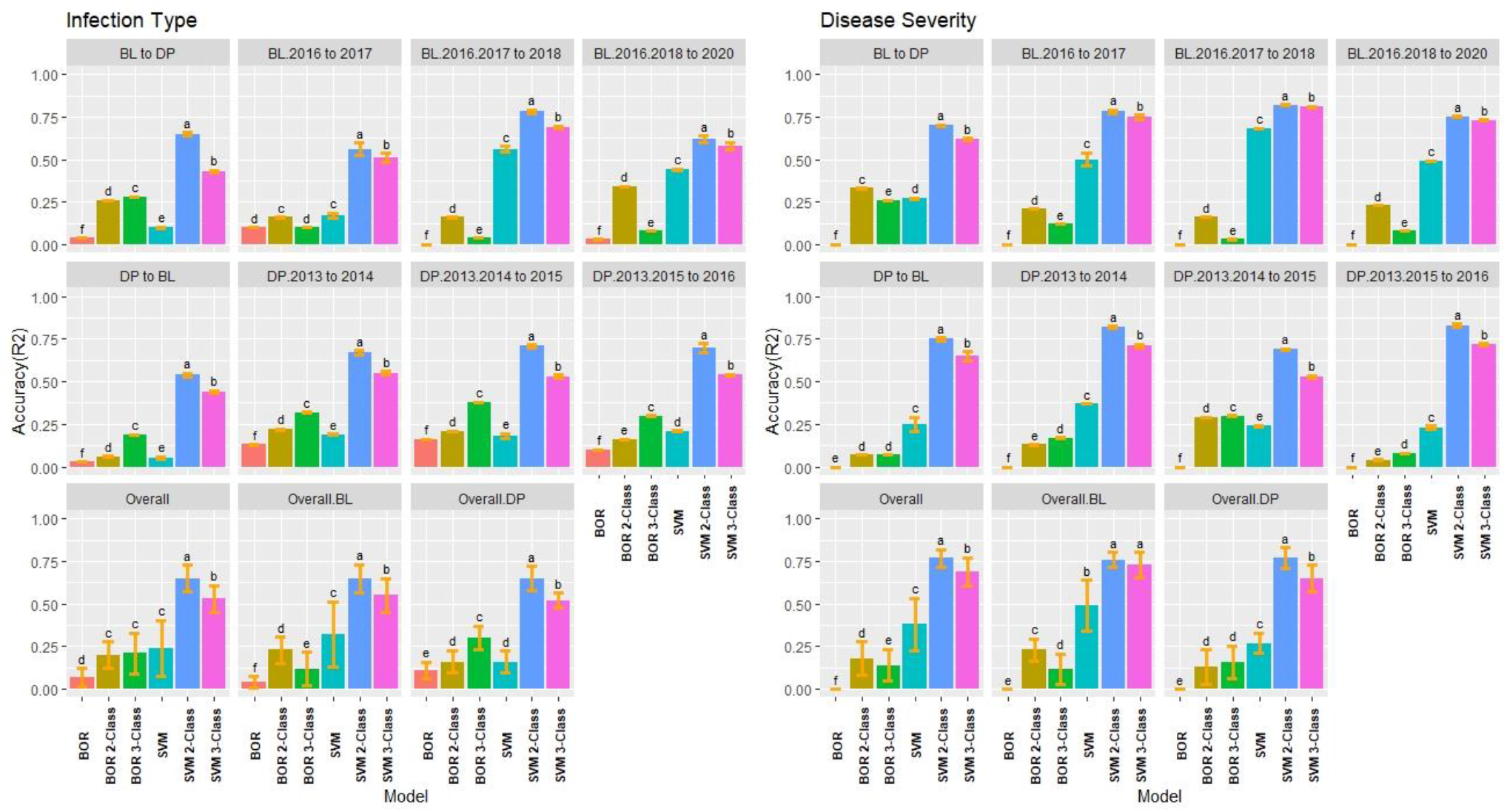
Pairwise comparisons of genomic selection classification model accuracy (*R*^*2*^) using validation sets for stripe rust infection type and disease severity. Pacific Northwest winter wheat diversity panel (DP) lines phenotyped from 2013 to 2016 in Central Ferry (DP_CF) and Pullman (DP_P), WA. Washington State University breeding lines phenotyped from 2016 to 2020 in Lind (BL_L) and Pullman (BP_L), WA. Model comparison across the DP (Overall.DP), BL (Overall.BL), and Overall scenarios. Models labeled with the same letter are not significantly different. Bars indicate standard errors.

SEV displayed similar results with IT, but the BOR model had zero *r* for all scenarios. However, the accuracies increased in the BOR with the reduced class scales (Figure 7). The 2 and 3-class SVM models displayed very high accuracy with 2-Class SVM reaching an overall class accuracy of 0.83 and maintained the high accuracy predicting the other population for both the BL and DP validation scenarios. Combining years did not result in improved accuracy. Further, the kappa values were very low except for the 2 and 3-Class SVM models reaching kappa values of 0.63 in the DP (Figure S11).

### 3.8 Validation set relative efficiency

The RE of the regression models were high in the validation scenarios reaching RE values of 0.85 using the SQRT rrBLUP model (Figure 8). The BOR and SVM models displayed relatively low RE values compared to the regression models. This was presumably due to the BOR not being able to predict the phenotypic values of the majority of the lines; however, the RE was higher for the classification models than the cross-validations with only one scenario having a negative value (−0.20). For overall comparisons, there were significant differences compared to the cross-validation scenarios. The SQRT rrBLUP reached the highest overall RE with 0.60. Further, the RE values were higher in the DP than the BL. Combining years was related to an increased RE for the regression models.

**Figure 8.**
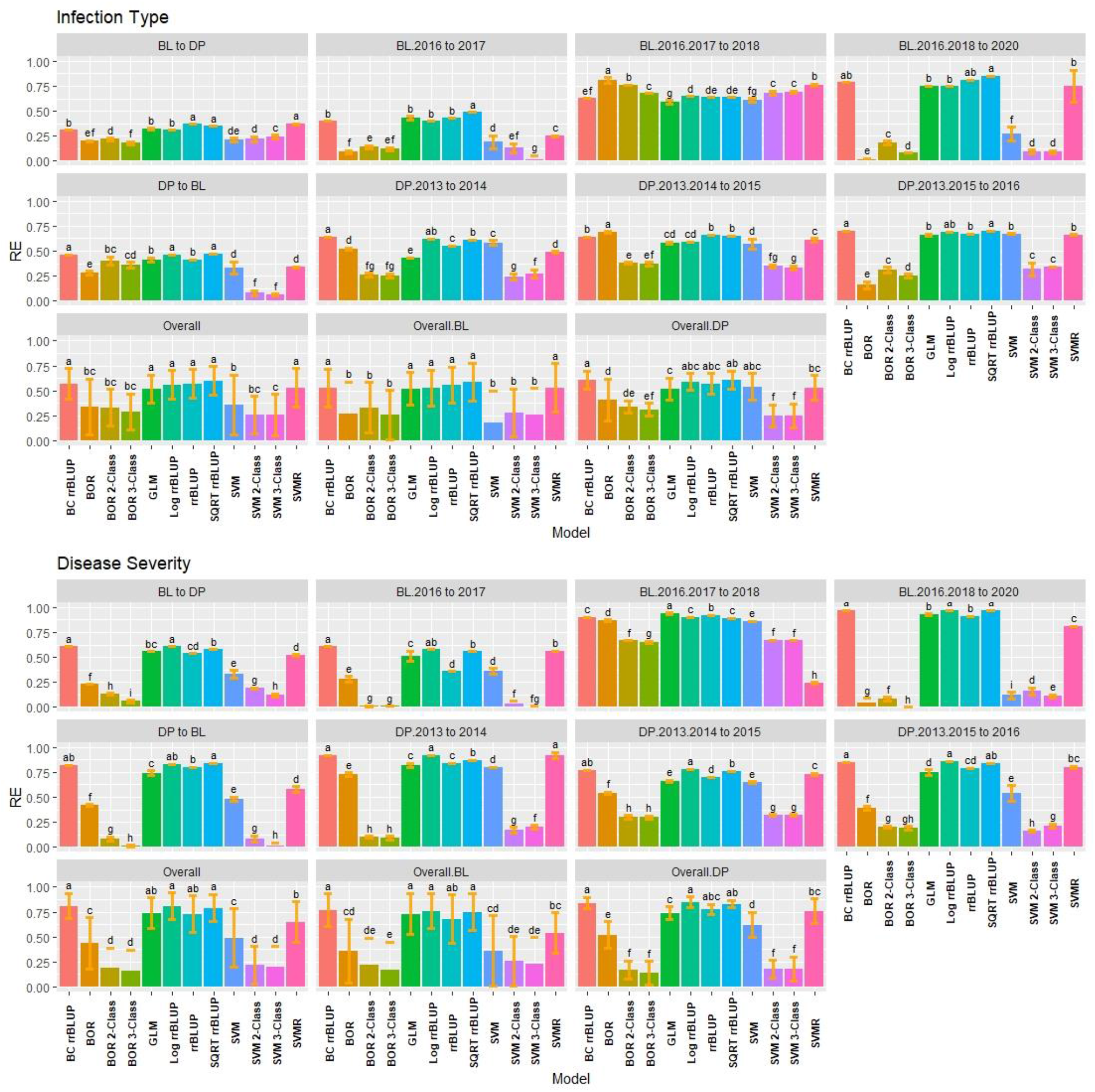
Pairwise comparisons of genomic selection regression and classification model relative efficiency (RE) using validation sets for stripe rust infection type. Pacific Northwest winter wheat diversity panel (DP) lines phenotyped from 2013 to 2016 in Central Ferry (DP_CF) and Pullman (DP_P), WA. Washington State University breeding lines phenotyped from 2016 to 2020 in Lind (BL_L) and Pullman (BL_P), WA. Model comparison across the DP (Overall.DP), BL (Overall.BL), and Overall scenarios. Models labeled with the same letter are not significantly different. Bars indicate standard errors.

Consistent trends for SEV were observed with the transformed rrBLUP models RE values of 0.97, displaying very high RE compared to phenotypic selection (Figure S12). The RE for SEV were relatively high for the rrBLUP and SVMR models predicting into the other population using the DP as the training population ranging from 0.58 to 0.86. Further displaying the ability for the regression models to accurately predict across years and populations while dealing with skewed phenotypes.

## 4 Discussion

Genomic selection has many advantages over traditional phenotypic selection and marker-assisted selection. Increased genetic gain and improved trait selection can be achieved by using GS (Heffner et al., 2010; Rutkoski et al., 2015; Michel et al., 2017). Furthermore, GS can aid in selection for traits dependent on the environment to display variation especially in years with little to no phenotypic variation for phenotypic selection. Plant breeding programs continually select and improve disease resistance due to the evolving race and pathogen changes along with the breakdown of resistance genes. Due to the high levels of resistance targeted within most plant breeding programs, positively skewed phenotypes generally result when selecting for disease resistance. Furthermore, disease resistance is commonly phenotyped in ordinal scales and percentages. The skewed and ordinal phenotypes pose challenges to utilizing regression models for genomic selection (Montesinos-López et al., 2015a). However, most GS studies treat disease resistance as continuous values and utilize regression models and transformations for prediction, while only a few studies have used classification methods (Ornella et al., 2012, 2014; Rutkoski et al., 2014; Arruda et al., 2016; Muleta et al., 2017; González-Camacho et al., 2018; Merrick et al., 2021). In the current study, we compared several regression and classification methods for genomic prediction for skewed phenotypes in the context of stripe rust resistance in winter wheat and identified the best approaches to use for predicting traits with skewed distributions.

When utilizing GS for resistance to diseases such as stripe rust, GS approaches can capture the additive effects of APR and are therefore relevant for accumulating favorable alleles for rust resistance. GS can reach high levels of accuracy for stripe rust and other rust diseases (Ornella et al., 2012; Rutkoski et al., 2014, 2015; Muleta et al., 2017; Merrick et al., 2021). Because of the high levels of resistance and high heritability of disease resistance in most breeding programs, phenotypic selection and marker-assisted selection have been shown to be successful (Lande and Thompson, 1990). Even so, GS has been shown to be superior to marker-assisted selection in selecting for APR in the presence of major resistance genes (Merrick et al., 2021).

### 4.1 Accuracy of regression models

Regression models assume continuous and normally distributed phenotypes (Montesinos-López et al., 2015c). In the current study, the BL and DP populations displayed skewed distributions for both IT and SEV with inflations of zero due to the high levels of disease resistance. Among the primary approaches used for phenotypes that do not follow a normal distribution are disregarding the lack of normality or transforming the phenotypes to a normal distribution (Montesinos-López et al., 2015b). In the current study, we observed that even with the skewed distributions, the rrBLUP model without transformed phenotypes still displayed high accuracies and performed similarly to the highest performing SQRT rrBLUP model in many scenarios. For example, there were no significant differences between SQRT rrBLUP and rrBLUP in the overall comparisons in the cross-validation (Figure 3) or validation set scenarios (Figures 6 and 7). These results support previous studies that utilized rrBLUP models for disease resistance (Rutkoski et al., 2014, 2015; Juliana et al., 2017; Muleta et al., 2017; Merrick et al., 2021). The performance of the untransformed rrBLUP model may be due to the central limit theorem which argues that given a sufficient number of observations, the sampling distribution of the means can be assumed to be approximately normal (Stroup, 2015).

Transformations were introduced to stabilize variance and fulfill the homogenous variance assumption of linear regression models (Bartlett, 1947). However, transformations have shown to produce a loss of accuracy and power in small sample size (Stroup, 2015). Further, in our study, the log and BC transformations displayed lower accuracy than the SQRT transformation. One of the problems with log transformations is the large number of zeros due to the presence of highly resistant lines in both the BL and DP populations. This occurrence constrains the transformation to stabilize variance and transform the phenotypes to follow a normal distribution (O’Hara and Kotze, 2010). Further, log transformations yield downwardly biased estimates, whereas SQRT does not (Stroup, 2015). The BC transformations is a powerful transformation that raise numbers to an exponent; nonetheless, BC requires lambda estimation and can theoretically be the same as the SQRT transformation at λ= 0.50 (Osborne, 2019). Therefore, if the optimal λ is not chosen correctly, the BC may not appropriately stabilize the variance of the data.

The SQRT transformation proved to perform very well for both accuracy and RE across populations, cross-validations, and validation scenarios in the current study. The SQRT transformations showed the ability to have higher accuracy and reduced RMSE compared with the untransformed data for the rrBLUP model. In Poisson distributions similar to the skewed phenotypes of our study, the variance is equal to the mean, and the SQRT is recommended to stabilize variance in those scenarios (Bartlett 1947); this could have resulted to increased performance for the SQRT transformation. Overall, the appropriate method must be chosen carefully when implementing data transformation on breeding programs.

Using the GLM model, high accuracy (0.66 and 0.76) in both the DP and BL training populations were observed. The performance of GLM was noted to be dependent on the distribution of the phenotypes. The GLM performed similarly to the rrBLUP model in the highly skewed selected BL population, but displayed statistically significant lower accuracies in the less skewed unselected DP population. Poisson GLMs, which were implemented in the present work, have been shown to display superior accuracy while correctly fitting the data (O’Hara and Kotze, 2010; Montesinos-López et al., 2015b, 2016, 2020; Stroup, 2015). The Poisson GLM accurately models count and ordinal data and is therefore suited for skewed phenotypes such as disease resistance (Ornella et al., 2014; Montesinos-López et al., 2015a, 2016). Further, the GLM models outperformed deep learning models in a previous study (Montesinos-López et al., 2020). The utilization of GLMs should be implemented in scenarios with the appropriate distribution of phenotypes.

Non-parametric models such as SVMR, which has no underlying assumption on the distribution of the phenotypes, performed better than the LOG and BC transformations, and similar to the GLM model in the current study. Previously, SVMR model has been shown to have superior prediction and RE values over parametric and semi-parametric models for predicting disease resistance due to the skewed phenotypes (González-Camacho et al., 2018). This demonstrates that the SVMR can accurately predict skewed phenotypes without the need to transform the data. SVM regression maps samples from a predictor space to a high-dimensional feature space using a non-linear kernel function and then completes linear regression in the feature space (Jannink et al., 2010). Consequently, this creates the ability for the SVMR to predict skewed phenotypes and allows the model to learn the complexity of the training population without imposing structure on the data (González-Camacho et al., 2018).

The SQRT rrBLUP models performed better than the SVMR model in overall prediction accuracy across many scenarios. The lack of advantage in regression scenarios was also observed by Ornella et al. (2014), where reproducing kernel Hilbert Space models were observed to be statistically significant for all yield data sets over SVM and random forest models. In the current study, the subordinate performance of the SVMR models is presumably due to the mostly additive effect of stripe rust resistance. Once the skewed phenotypes are properly modeled, the use of non-parametric models that also model non-additive effects disappears (Ornella et al., 2014; Poland and Rutkoski, 2016).

### 4.2 Accuracy of classification models

In the present study, Bayesian Ordinal Regression (BOR) models displayed the lowest accuracies and RE across all the classification and regression models, particularly in the DP population. Conversely, the BOR models using reduced classes reached the highest overall class accuracy over all models with *r*= 0.99 for the BL. In contrast, when the accuracy was high in the BL, the kappa values were low. The opposite was shown in the DP with low overall class accuracies and moderate kappa values indicating that the high overall class accuracy and low kappa values was a result of the BOR model consistently predicting zeros and the inability to predict the other classes. Further, in the validation sets, the BOR performed very poorly, and resulted in near zero overall class accuracy and kappa values for both IT and SEV. The BOR model uses ordinal regression that is suitable for count and censored data; nevertheless, the BOR model uses the probit link function that does not explicitly model non-normal distribution such as the Poisson distribution model by the GLM model in our study (Montesinos-López et al., 2015a). Altogether, our results showed that the BOR model is not appropriate for the highly skewed phenotypes in our study.

We also used SVMs as the non-parametric machine learning model for both regression and classification. The advantage in classification and regression using SVM models for disease resistance has been previously demonstrated (Ornella et al., 2014; González-Camacho et al., 2018). Contrary to the BOR results, the SVMs consistently displayed high accuracies throughout the locations and years for both DP and BL training populations. However, the SVM showed lower kappa values than the BOR in many scenarios. This was not the trend in the validation sets, where the full scale BOR and SVM displayed poor accuracy and kappa values. The consistent accuracy of SVM over BOR may be due to the non-parametric nature of the SVM models. The SVM model is implemented similar to the SVMR model and use soft classifiers to calculate the probability of the class rather than hard classifiers which directly targets the decision boundary and allows the model to be flexible (Ornella et al., 2014). Based on the results for BOR and SVM, classification models need to be compared by both overall class accuracy as well as a metric such as kappa that accounts for individual class accuracy.

The precision of the classification models depends on the number of individuals in a given class. In our study, we implemented up-sampling (i.e., random sampling with replacement) to increase the minority class to the same size of the majority class and reduce the effect of class imbalance (Kuhn, 2008). However, our results showed that with imbalanced class frequency due to skewed phenotypes, even resampling techniques such as up-sampling failed to accurately predict disease resistance. Another approach to deal with class frequency is to reduce the number of overall classes. We then binned classes to create a 2 and 3-class prediction scenarios. Reducing the class scale to two creates a binary classification model which has been shown to outperform other regression and classification models (Ornella et al., 2014). By reducing the number of classes, we also decreased the effect of class imbalances. Accuracy as well as kappa increased specifically for the SVM by reducing the class scales. This observation was seen even in the validation sets, which resulted in the SVM 2-class models achieving both high accuracy and kappa values, consistent with previous studies on the effects of reduced classes (Ornella et al., 2014; González-Camacho et al., 2016). Therefore, by reducing the class scale, classification models such as SVM can accurately predict skewed phenotypes such as disease resistance.

### 4.3 Relative efficiency

Relative efficiency compares the expected genetic gain when selecting based on GEBVs compared to phenotypic selection. The RE can be used as an indicator of the performance of a model when used for truncation selection and expected genetic gain (Ornella et al., 2014; González-Camacho et al., 2018). Since classification and regression do not use the same metrics for performance, simply comparing accuracies is not possible; hence, we used RE for comparisons. A selection intensity of 15% was used based on a previous study (Ornella et al., 2014). In the current work, the rrBLUP models and SMVR displayed high RE values across both cross-validation and validation sets for IT and SEV with values above 0.90. SVMR models have been shown to have superior RE values for classification in disease resistance (Ornella et al., 2014). The high RE values indicated that accuracy is linear in the regression models, but this was not the case for the classification models. The classification models displayed relatively low RE values and in some cases negative values. Both the SVM and BOR models displayed the inability to select the top 15% performers for stripe rust resistance. The large amounts of zeros (i.e., disease resistant phenotypes) skew the prediction accuracy for the classification models to the very high, with low kappa and RE values. The classification models failed to overcome the skewed phenotypes even with up-sampling and reduction of classes. Therefore, similar to our results for prediction accuracy, regression models outperformed classification models and displayed their ability to predict and select skewed phenotypes.

### 4.4 Training Population Comparison

We compared the performance of GS models in different training populations, environments, and phenotypic distributions. The effect of environment was less apparent than the effect of distribution. The differences in distribution of phenotypes for disease resistance is readily apparent between populations. The two populations were used to compare the effects of a selected and unselected population with varying degrees and sources of resistance. The BL population, consisting of WSU breeding lines that were selected for disease resistance prior to field trials, is extremely skewed for both IT and SEV. Therefore, there is already a selection pressure for high levels of resistance to stripe rust in the current study. In contrast, the DP appears less skewed with more variation for disease resistance; a consequence of the population consisting of diverse varieties from multiple breeding programs in the Pacific Northwest region of the US. The DP included lines from the WSU breeding program, but the other varieties were not bred and selected specifically for resistance to the stripe rust races present in our study. Additionally, the sources of stripe rust resistance genes vary more in the DP compared to the BL. The frequency and type of stripe rust races along with major genes for stripe rust resistance for these two populations were compared in depth in Merrick et al. (2021).

The differences in skewness between the populations affected the performance of the GS models in each population. Since the GLM models the Poisson distribution, it accurately predicted the extremely skewed BL trials similar to the other regression models; however, it displayed lower accuracies in the DP. In addition to the distribution that is modeled, the skewness affects the frequency of classes used in classification models. In the extremely skewed BL, the classification models have high accuracy and low kappa, displaying the prediction of mainly zeros. However, as mentioned previously, the reduction of classes helps decrease the effect of class imbalance and increased accuracy. The differences in accuracies between populations can also be attributed to the genetic relatedness of the populations (Asoro et al., 2011). The effect of the population on accuracy is due to both population structure and genetic relatedness (Habier et al., 2007; Asoro et al., 2011; Mirdita et al., 2015). We used the elbow method to determine the number of clusters when examining PCs for our populations and resulted in four distinct clusters. Consequently, the prediction accuracy for the BL cross-validations were higher than the DP. When independently predicting other populations as seen in the validations sets, we generally observe a decrease in accuracy (Merrick and Carter, 2021; Merrick et al., 2021). Interestingly though, an increased in accuracy with the BL predicting the DP in the validation sets, with the opposite observation when predicting the BL with the DP was observed. However, this was only seen in the regression models for predicting SEV in the validation sets. Further, this trend is not seen in the classification models which display consistent accuracy across validation scenarios. This may be due to the effect of predicting a less skewed population which regression models generally have better performance compared to predicting more skewed distributions (Montesinos-López et al., 2015a).

The increase in prediction accuracy with the increased combination of years in both our cross-validation and validation sets can be attributed to the increase of phenotypic data points and decrease in skewness and accounting for the genotype-by-environment interaction (GEI). The trials in our study were dependent on the natural occurrence and pressure of stripe rust. Therefore, the skewness of the populations, individual years, and locations may not only be due the levels of resistance within the populations, but also due to the general disease pressure for stripe rust. By combining environments, we can account for the GEI in our phenotypic adjustments and increase our prediction accuracy (Jarquín et al. 2014; Crossa et al. 2014; Haile et al. 2020; Merrick and Carter 2021; Merrick et al. 2021). The increased accuracy by accounting for GEI can be seen in the validation sets. The DP displayed higher accuracies in the validation sets as the DP comprised of the same lines each year, whereas the BL consists of different lines in both years and locations. By screening the same lines each year, the environmental effect can be effectively accounted for. However, the trend for increasing accuracy and RE values by combining years was not seen in the classification models. This was due to the continued large class imbalances even when combining years. Therefore, there is a need to develop training populations carefully to balance class frequencies for the classification models. Even so, the reduced class SVM models displayed the ability to overcome the class frequencies regardless of year combinations. Overall, the rrBLUP and reduced class classification models displayed the ability to accurately predict populations and environments with skewed phenotypes.

### 4.5 Applications in Breeding

Genomic selection is becoming more cost-effective due to the decreasing costs of high-throughput genotyping. With the increased use of GS comes its utilization for the prediction of complex traits (e.g., disease resistance) which do not always follow the assumptions of the commonly used models (Montesinos-López et al., 2015a). Instead of applying the same approach to every trait, breeders will need to customize their GS models to achieve accurate GEBVs for selection. With the integration of data science and plant breeding, the availability of different prediction models has resulted in an increased efficiency of implementing GS for a wide range of traits. This study showed that with the appropriate choice of model and transformation, even the commonly used GS regression model, rrBLUP, can be utilized for predicting complex traits, such as stripe rust resistance, that do not follow a normal distribution. Further, this study demonstrated the ability to integrate selection decisions and GS by utilizing classification models. Reducing classes resulted in higher predictions due to decreasing the number of outcomes the models need to account for, especially for classes with only a few observations. Moreover, by reducing the number of classes, we not only predict resistance more accurately, but also couple in selection decisions. By reducing the number of classes for IT from ten to two, we can either keep or discard lines. Ultimately, by using various GS schemes with regression and classification models, breeders can reduce the number of selection decisions made for disease resistance and focus on selecting other important traits such as grain yield.

## 5 Conclusions

This study compared GS regression and classification models’ ability to accurately predict populations with different levels of disease resistance and distributions. The varying results for the classification and transformation methods displayed the need to choose the prediction model carefully based on the phenotype distribution. For trials that display a Poisson distribution that is skewed to lower ordinal values, a GLM or reduced class binomial classification model can be implemented. However, the SQRT and SVMR models displayed the flexibility across varying distributions, and consistently predicted stripe rust with high accuracies. Moreover, combining years increased the prediction accuracies for regression models, but failed to increase the overall class accuracy for classification models due to imbalance class frequencies. Additionally, regression models displayed high RE indicating their ability to select accurately like phenotypic selection. Overall, SQRT transformation using rrBLUP and SVM regression models displayed the highest combination of accuracy and relative efficiency across the regression and classification models. Further, a classification system based on SVM with a 2-Class scale can be implemented not only to predict resistance more accurately, but also couple in selection decisions. This study showed that breeders can use linear and non-parametric regression models using their own breeding lines over combined years to accurately predict skewed phenotypes.

## Supporting information

Supplementary tables are available in a separate file on BioRvx.

## 7 Author Contributions

LFM: conceptualized the idea, analyzed data, and drafted the manuscript; DNL: reviewed and edited the manuscript; XC: reviewed and edited the manuscript; AHC: supervised the study, conducted field trials, edited the manuscript, and obtained the funding for the project.

## 9 Acknowledgments

The authors would like to acknowledge the Washington State University Winter Wheat Breeding Program personnel Gary Shelton and Kyall Hagemeyer for plot maintenance and data collection under field conditions. We would also like to thank Adrienne Burke, Gina Brown-Guedira, Jared Smith, Brian Ward, and staff at the Eastern Regional Small Grains Genotyping Laboratory for their assistance with DNA library prep and GBS sequencing and analysis.

## 10 Data Availability Statement

The datasets generated for this study can be found at https://github.com/lfmerrick21/Regression-vs-Classification.

## 11 Supplementary Material

Supplementary tables are available in a separate file on BioRvx.

## References

Appels, R., Eversole, K., Stein, N., Feuillet, C., Keller, B., Rogers, J., et al. (2018). Shifting the limits in wheat research and breeding using a fully annotated reference genome. Science 361, eaar7191. doi:10.1126/science.aar7191.

Arruda, M. P., Lipka, A. E., Brown, P. J., Krill, A. M., Thurber, C., Brown-Guedira, G., et al. (2016). Comparing genomic selection and marker-assisted selection for Fusarium head blight resistance in wheat (Triticum aestivum L.). Mol. Breed. 36, 84. doi:10.1007/s11032-016-0508-5.

Asoro, F. G., Newell, M. A., Beavis, W. D., Scott, M. P., and Jannink, J.-L. (2011). Accuracy and training population design for genomic selection on quantitative traits in elite North American oats. Plant Genome J. 4, 132. doi:10.3835/plantgenome2011.02.0007.

Bartlett, M. S. (1947). The use of transformations. Biometrics 3, 39–52. doi:10.2307/3001536.

Bradbury, P. J., Zhang, Z., Kroon, D. E., Casstevens, T. M., Ramdoss, Y., and Buckler, E. S. (2007). TASSEL: software for association mapping of complex traits in diverse samples. Bioinformatics 23, 2633–2635.

Browning, B. L., Zhou, Y., and Browning, S. R. (2018). A one-penny imputed genome from next-generation reference panels. Am. J. Hum. Genet. 103, 338–348. doi:https://doi.org/10.1016/j.ajhg.2018.07.015.

Chen, X. (2013). High-temperature adult-plant resistance, key for sustainable control of stripe rust. Am. J. Plant Sci. 4, 608–627. https://doi.10.4236/ajps.2013.43080

Chen, X. (2020). Pathogens which threaten food security: Puccinia striiformis, the wheat stripe rust pathogen. Food Security 12, 239–251. https://doi.org/10.1007/s12571-020-01016-z

Chen, X. and Line, R. F. (1995a). Gene action in wheat cultivars for durable, high-temperature, adult-plant resistance and interaction with race-specific, seedling resistance to Puccinia striiformis. Phytopathology 85, 567–572.

Chen, X. and Line, R. F. (1995b). Gene number and heritability of wheat cultivars with durable, high-temperature, adult-plant (HTAP) resistance and interaction of HTAP and race-specific seedling resistance to Puccinia striiformis. Phytopathology 85, 573–578.

Crossa, J., Pérez, P., Hickey, J., Burgueño, J., Ornella, L., Cerón-Rojas, J., et al. (2014). Genomic prediction in CIMMYT maize and wheat breeding programs. Heredity 112, 48–60. doi:10.1038/hdy.2013.16.

Cullis, B. R., Smith, A. B., and Coombes, N. E. (2006). On the design of early generation variety trials with correlated data. J. Agric. Biol. Environ. Stat. 11, 381. doi:10.1198/108571106X154443.

de Mendiburu, F., and de Mendiburu, M. F. (2019). Package ‘agricolae.’ R Package Version, 1.2-8.

Elshire, R. J., Glaubitz, J. C., Sun, Q., Poland, J. A., Kawamoto, K., Buckler, E. S., et al. (2011). A robust, simple genotyping-by-sequencing (GBS) approach for high diversity species. PLoS ONE 6, e19379. doi:10.1371/journal.pone.0019379.

Endelman, J. B. (2011). Ridge regression and other kernels for genomic selection with R package rrBLUP. Plant Genome J. 4, 250. doi:10.3835/plantgenome2011.08.0024.

Federer, W. F. (1956). Experimental design. LWW.

Gareth, J., Daniela, W., Trevor, H., and Robert, T. (2013). An introduction to statistical learning: with applications in R. Spinger.

Glaubitz, J. C., Casstevens, T. M., Lu, F., Harriman, J., Elshire, R. J., Sun, Q., et al. (2014). TASSEL-GBS: A High capacity genotyping by sequencing analysis pipeline. PLOS ONE 9, e90346. doi:10.1371/journal.pone.0090346.

Goldman, I. (2019). Plant Breeding Reviews. John Wiley & Sons.

González-Camacho, J. M., Crossa, J., Pérez-Rodríguez, P., Ornella, L., and Gianola, D. (2016). Genome-enabled prediction using probabilistic neural network classifiers. BMC Genomics 17, 1–16.

González-Camacho, J. M., Ornella, L., Pérez-Rodríguez, P., Gianola, D., Dreisigacker, S., and Crossa, J. (2018). Applications of machine learning methods to genomic selection in breeding wheat for rust resistance. Plant Genome 11, 170104. doi:10.3835/plantgenome2017.11.0104.

Habier, D., Fernando, R. L., and Dekkers, J. C. M. (2007). The impact of genetic relationship information on genome-assisted breeding values. Genetics 177, 2389–2397. doi:10.1534/genetics.107.081190.

Haile, T. A., Walkowiak, S., N’Diaye, A., Clarke, J. M., Hucl, P. J., Cuthbert, R. D., et al. (2020). Genomic prediction of agronomic traits in wheat using different models and cross-validation designs. Theor. Appl. Genet. 134, 381–398. doi:https://doi.org/10.1007/s00122-020-03703-z.

Hastie, T., Qian, J., and Tay, K. (2016). An Introduction to glmnet.

Heffner, E. L., Lorenz, A. J., Jannink, J.-L., and Sorrells, M. E. (2010). Plant breeding with genomic selection: Gain per unit time and cost. Crop Sci. 50, 1681. doi:10.2135/cropsci2009.11.0662.

Hyndman, R. J., and Khandakar, Y. (2008). Automatic time series forecasting: the forecast package for R. J. Stat. Softw. 27, 1–22.

Jannink, J.-L., Lorenz, A. J., and Iwata, H. (2010). Genomic selection in plant breeding: from theory to practice. Brief. Funct. Genomics 9, 166–177. doi:10.1093/bfgp/elq001.

Jarquín, D., Crossa, J., Lacaze, X., Du Cheyron, P., Daucourt, J., Lorgeou, J., et al. (2014). A reaction norm model for genomic selection using high-dimensional genomic and environmental data. Theor. Appl. Genet. 127, 595–607. doi:http://dx.doi.org/10.1007/s00122-013-2243-1.

Juliana, P., Singh, R. P., Singh, P. K., Crossa, J., Huerta-Espino, J., Lan, C., et al. (2017). Genomic and pedigree-based prediction for leaf, stem, and stripe rust resistance in wheat. Theor. Appl. Genet. 130, 1415–1430. doi:10.1007/s00122-017-2897-1.

Kamiak (2021). High Performance Computing | Washington State University. High Perform. Comput. Available at: https://hpc.wsu.edu/ [Accessed January 21, 2021].

Karatzoglou, A., Smola, A., Hornik, K., and Karatzoglou, M. A. (2019). Package ‘kernlab.’ CRAN R Proj.

Kassambara, A., and Kassambara, M. A. (2020). Package ‘ggpubr.’

Klarquist, E., Chen, X., and Carter, A. (2016). Novel QTL for stripe rust resistance on chromosomes 4A and 6B in soft white winter wheat cultivars. Agronomy 6, 4. doi:10.3390/agronomy6010004.

Kuhn, M. (2008). Building predictive models in R using the caret package. J. Stat. Softw. 28, 1–26. doi:10.18637/jss.v028.i05.

Lande, R., and Thompson, R. (1990). Efficiency of marker-assisted selectionin the improvement of quantitative eraits. Genetics 124, 743–756.

Li, H., and Durbin, R. (2009). Fast and accurate short read alignment with Burrows–Wheeler transform. Bioinformatics 25, 1754–1760. doi:https://doi.org/10.1093/bioinformatics/btp324.

Line, R. F., and Qayoum, A. (1992). Virulence, aggressiveness, evolution and distribution of races of Puccinia striiformis (the cause of stripe rust of wheat) in North America, 1968-87. Tech. Bull. USA. Available at: http://agris.fao.org/agris-search/search.do?recordID=US9304750 [Accessed January 16, 2020].

Liu, L., Yuan, C. Y., Wang, M. N., See, D. R., Zemetra, R. S., and Chen, X. M. (2019). QTL analysis of durable stripe rust resistance in the North American winter wheat cultivar Skiles. Theor. Appl. Genet. 132, 1677–1691. doi:10.1007/s00122-019-03307-2.

Liu, Y., Qie, Y., Li, X., Wang, M., and Chen, X. (2020). Genome-wide mapping of quantitative trait loci conferring all-stage and high-temperature adult-plant resistance to stripe rust in spring wheat landrace PI 181410. Int. J. Mol. Sci. 21, 478. doi:10.3390/ijms21020478.

Merrick, L. F., Burke, A. B., Chen, X., and Carter, A. H. (2021). Breeding with major and minor genes: Genomic selection for quantitative disease resistance. Front. Plant Sci. 12, 1599. doi:10.3389/fpls.2021.713667.

Merrick, L. F., and Carter, A. H. (2021). Comparison of genomic selection models for exploring predictive ability of complex traits in breeding programs. Plant Genome 14, e20158. doi:10.1002/tpg2.20158.

Meyer, D., Dimitriadou, E., Hornik, K., Weingessel, A., Leisch, F., Chang, C.-C., et al. (2019). Package ‘e1071.’ R J.

Michel, S., Ametz, C., Gungor, H., Akgöl, B., Epure, D., Grausgruber, H., et al. (2017). Genomic assisted selection for enhancing line breeding: merging genomic and phenotypic selection in winter wheat breeding programs with preliminary yield trials. Theor. Appl. Genet. 130, 363–376. doi:10.1007/s00122-016-2818-8.

Mirdita, V., He, S., Zhao, Y., Korzun, V., Bothe, R., Ebmeyer, E., et al. (2015). Potential and limits of whole genome prediction of resistance to Fusarium head blight and Septoria tritici blotch in a vast Central European elite winter wheat population. Theor. Appl. Genet. 128, 2471–2481. doi:https://doi.org/10.1007/s00122-015-2602-1.

Montesinos-López, A., Montesinos-López, O. A., Crossa, J., Burgueño, J., Eskridge, K. M., Falconi-Castillo, E., et al. (2016). Genomic Bayesian prediction model for count data with genotype × environment interaction. G3amp58 GenesGenomesGenetics 6, 1165–1177. doi:10.1534/g3.116.028118.

Montesinos-López, O. A., Montesinos-López, A., Crossa, J., Burgueño, J., and Eskridge, K. (2015a). Genomic-enabled prediction of ordinal data with Bayesian logistic ordinal regression. G3 Genes Genomes Genetics 5, 2113–2126. doi:10.1534/g3.115.021154.

Montesinos-López, O. A., Montesinos-López, A., Pérez-Rodríguez, P., de los Campos, G., Eskridge, K., and Crossa, J. (2015b). Threshold models for genome-enabled prediction of ordinal categorical traits in plant breeding. G3 Genes Genomes Genetics 5, 291–300. doi:10.1534/g3.114.016188.

Montesinos-López, O. A., Montesinos-López, A., Pérez-Rodríguez, P., Eskridge, K., He, X., Juliana, P., et al. (2015c). Genomic prediction models for count data. J. Agric. Biol. Environ. Stat. 20, 533–554. doi:10.1007/s13253-015-0223-4.

Montesinos-López, O. A., Montesinos-López, J. C., Singh, P., Lozano-Ramirez, N., Barrón-López, A., Montesinos-López, A., et al. (2020). A multivariate Poisson deep learning model for genomic prediction of count data. G3 Genes Genomes Genet. 10, 4177–4190.

Muleta, K. T., Bulli, P., Zhang, Z., Chen, X., and Pumphrey, M. (2017). Unlocking Diversity in germplasm collections via genomic selection: A case study based on quantitative adult plant resistance to stripe rust in spring wheat. Plant Genome 10, 3. doi:10.3835/plantgenome2016.12.0124.

O’Hara, R. B., and Kotze, D. J. (2010). Do not log-transform count data: Do not log-transform count data. Methods Ecol. Evol. 1, 118–122. doi:10.1111/j.2041-210X.2010.00021.x.

Ornella, L., Pérez, P., Tapia, E., González-Camacho, J. M., Burgueño, J., Zhang, X., et al. (2014). Genomic-enabled prediction with classification algorithms. Heredity 112, 616–626. doi:10.1038/hdy.2013.144.

Ornella, L., Singh, S., Perez, P., Burgueño, J., Singh, R., Tapia, E., et al. (2012). Genomic prediction of genetic values for resistance to wheat rusts. Plant Genome 5, 136–148. doi:10.3835/plantgenome2012.07.0017.

Osborne, J. (2019). Improving your data transformations: Applying the Box-Cox transformation. Pract. Assess. Res. Eval. 15, 12. https://doi.org/10.7275/qbpc-gk17.

Pérez, P., and de los Campos, G. (2014). Genome-wide regression and prediction with the BGLR statistical package. Genetics 198, 483–495. doi:10.1534/genetics.114.164442.

Peterson RF, Campbell AB, Hannah AE (1948) A Diagrammatic scale for estimating rust intensity on leaves and stems of cereals. Can. J. Res. 26c, 496–500. https://doi.org/10.1139/cjr48c-033

Poland, J. A., and Rife, T. W. (2012). Genotyping-by-sequencing for plant breeding and genetics. Plant Genome J. 5, 92. doi:10.3835/plantgenome2012.05.0005.

Poland, J., and Rutkoski, J. (2016). Advances and challenges in genomic selection for disease resistance. Annu. Rev. Phytopathol. 54, 79–98. doi:10.1146/annurev-phyto-080615-100056.

R Core Team (2018). R: A Language and Environment for Statistical Computing. Vienna, Austria: R Foundation for Statistical Computing Available at: https://www.R-project.org/.

Riedelsheimer, C., Technow, F., and Melchinger, A. E. (2012). Comparison of whole-genome prediction models for traits with contrasting genetic architecture in a diversity panel of maize inbred lines. BMC Genomics 13, 452. doi:https://doi.org/10.1186/1471-2164-13-452.

Rutkoski, J. E., Poland, J. A., Singh, R. P., Huerta-Espino, J., Bhavani, S., Barbier, H., et al. (2014). Genomic selection for quantitative adult plant stem rust resistance in wheat. Plant Genome 7, 0. doi:10.3835/plantgenome2014.02.0006.

Rutkoski, J., Singh, R. P., Huerta-Espino, J., Bhavani, S., Poland, J., Jannink, J. L., et al. (2015). Efficient use of historical data for genomic selection: A case study of stem rust resistance in wheat. Plant Genome 8, 0. doi:10.3835/plantgenome2014.09.0046.

SAS Institute, Inc (2011). SAS® 9.3 system options: Reference. SAS Institute Inc Cary, NC.

Schmidt, P., Hartung, J., Bennewitz, J., and Piepho, H.-P. (2019). Heritability in plant breeding on a genotype-difference basis. Genetics 212, 991–1008.

Stroup, W. W. (2015). Rethinking the analysis of non-normal data in plant and soil science. Agron. J. 107, 811–827. doi:10.2134/agronj2013.0342.

Wang, X., Xu, Y., Hu, Z., and Xu, C. (2018). Genomic selection methods for crop improvement: Current status and prospects. Crop J. 6, 330–340. doi:10.1016/j.cj.2018.03.001.

Ward, B. P., Brown-Guedira, G., Tyagi, P., Kolb, F. L., Van Sanford, D. A., Sneller, C. H., et al. (2019). Multienvironment and multitrait genomic selection models in unbalanced early-generation wheat yield trials. Crop Sci. 59, 491. doi:10.2135/cropsci2018.03.0189.

Wickham, H. (2011). ggplot2. WIREs Comput. Stat. 3, 180–185. doi:10.1002/wics.147.

