## Supplementary tables are available in a separate file on BioRvx. for "Classification and Regression Models for Genomic Selection of Skewed Phenotypes: A Case for Disease Resistance in Winter Wheat (*Triticum aestivum* L.)"

### *Supplementary Material*

#### 1 Supplementary Figures

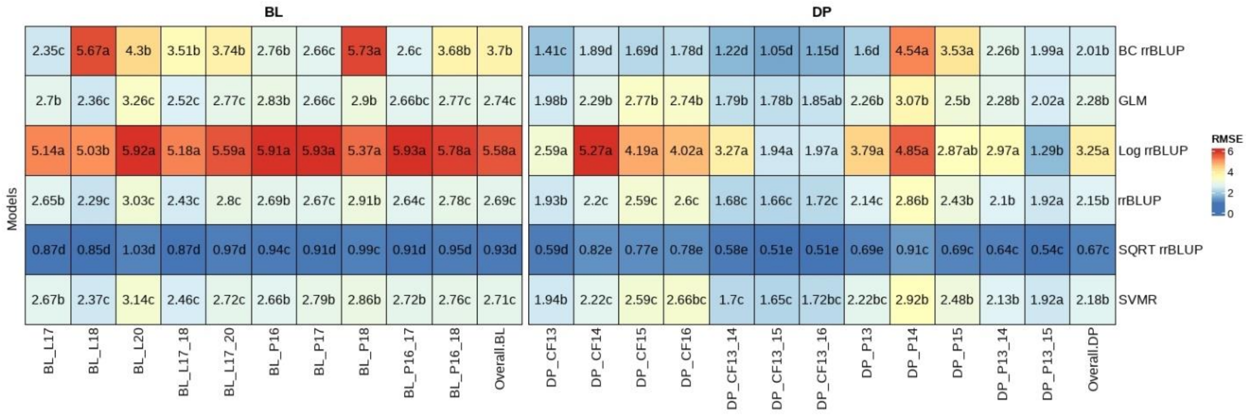


**Figure S1.** Heatmap of genomic selection regression model RMSE and pairwise comparisons using cross-validations for stripe rust infection type. Pacific Northwest winter wheat diversity panel (DP) lines phenotyped from 2013 to 2016 in Central Ferry (DP_CF) and Pullman (DP_P), WA. Washington State University breeding lines phenotyped from 2016 to 2020 in Lind (BL_L) and Pullman (BL_P), WA. Model comparison across the DP (Overall.DP), BL (Overall.BL), and Overall scenarios. Models labeled with the same letter are not significantly different (*P*-value =0.05).

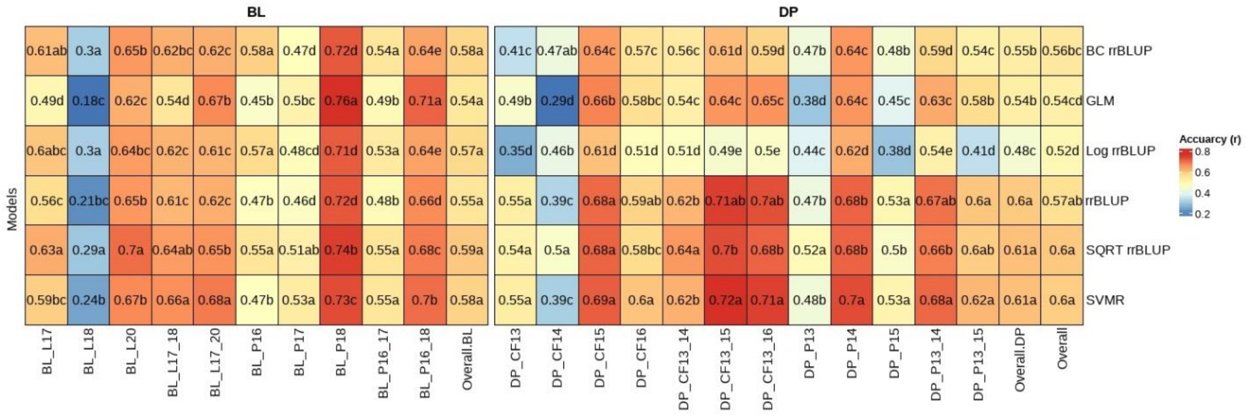


**Figure S2.** Heatmap of genomic selection regression model accuracy (*r*) and pairwise comparisons using cross-validations for stripe rust disease severity (SEV). Pacific Northwest winter wheat diversity panel (DP) lines phenotyped from 2013 to 2016 in Central Ferry (DP_CF) and Pullman (DP_P), WA. Washington State University breeding lines phenotyped from 2016 to 2020 in Lind (BP_L) and Pullman (BL_P), WA. Model comparison across the DP (Overall.DP), BL (Overall.BL), and Overall scenarios. Models labeled with the same letter are not significantly different (*P*-value =0.05).

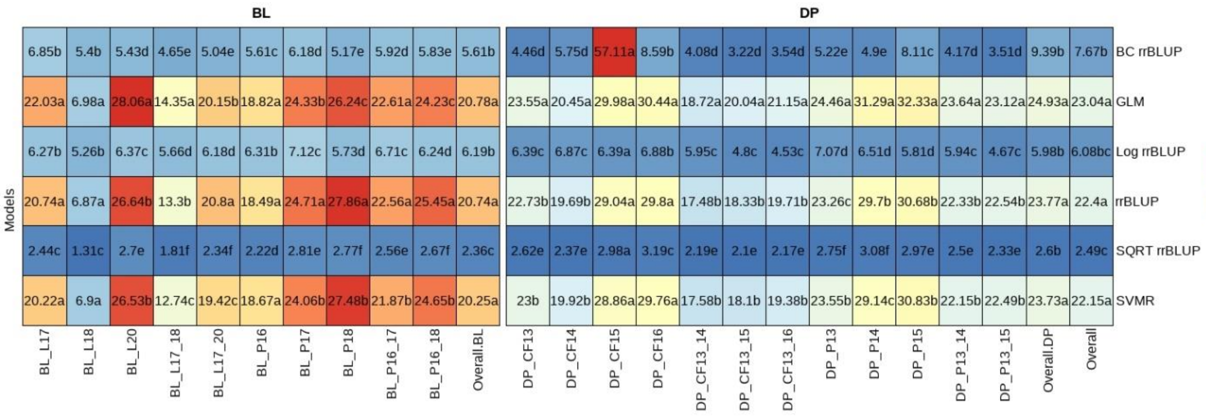


**Figure S3.** Heatmap of genomic selection regression model accuracy RMSE and pairwise comparisons using cross-validations for stripe rust disease severity. Pacific Northwest winter wheat diversity panel (DP) lines phenotyped from 2013 to 2016 in Central Ferry (DP_CF) and Pullman (DP_P), WA. Washington State University breeding lines phenotyped from 2016 to 2020 in Lind (BP_L) and Pullman (BL_P), WA. Model comparison across the DP (Overall.DP), BL (Overall.BL), and Overall scenarios. Models labeled with the same letter are not significantly different (*P*-value =0.05).

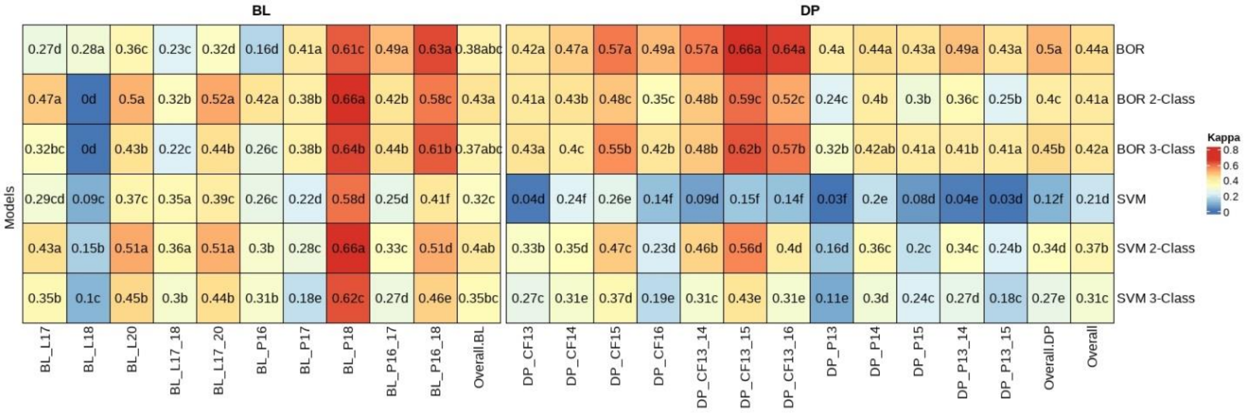


**Figure S4.** Heatmap of genomic selection classification model kappa and pairwise comparisons using cross-validations for stripe rust infection type. Pacific Northwest winter wheat diversity panel (DP) lines phenotyped from 2013 to 2016 in Central Ferry (DP_CF) and Pullman (DP_P), WA. Washington State University breeding lines phenotyped from 2016 to 2020 in Lind (BL_L) and Pullman (BL_P), WA. Model comparison across the DP (Overall.DP), BL (Overall.BL), and Overall scenarios. Models labeled with the same letter are not significantly different (*P*-value =0.05).

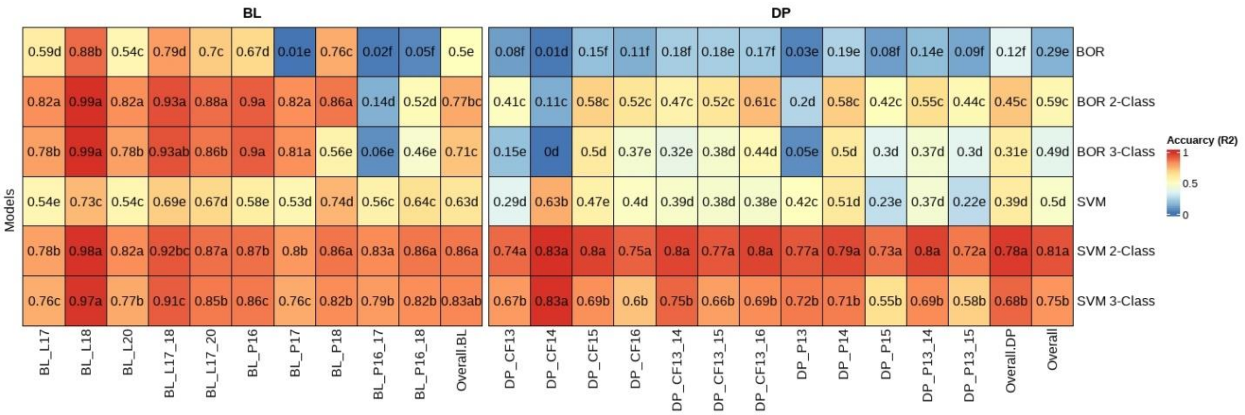


**Figure S5.** Heatmap of genomic selection classification model accuracy ($R^2$) and pairwise comparisons using cross-validations for stripe rust disease severity. Pacific Northwest winter wheat diversity panel (DP) lines phenotyped from 2013 to 2016 in Central Ferry (DP_CF) and Pullman (DP_P), WA. Washington State University breeding lines phenotyped from 2016 to 2020 in Lind (BL_L) and Pullman (BL_P), WA. Model comparison across the DP (Overall.DP), BL (Overall.BL), and Overall scenarios. Models labeled with the same letter are not significantly different (*P*-value =0.05).

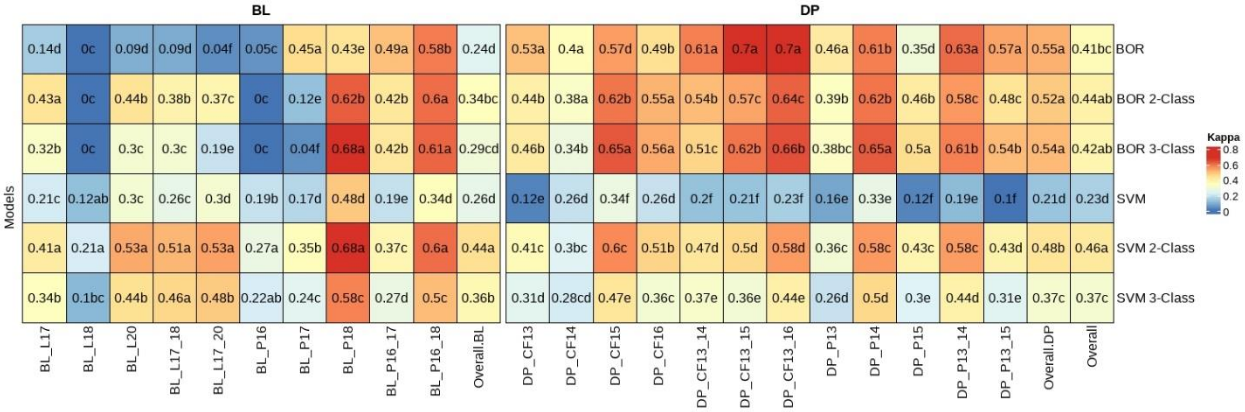


**Figure S6.** Heatmap of genomic selection classification model kappa and pairwise comparisons using cross-validations for stripe rust disease severity. Pacific Northwest winter wheat diversity panel (DP) lines phenotyped from 2013 to 2016 in Central Ferry (DP_CF) and Pullman (DP_P), WA. Washington State University breeding lines phenotyped from 2016 to 2020 in Lind (BL_L) and Pullman (BL_P), WA. Model comparison across the DP (Overall.DP), BL (Overall.BL), and Overall scenarios. Models labeled with the same letter are not significantly different (*P*-value =0.05).

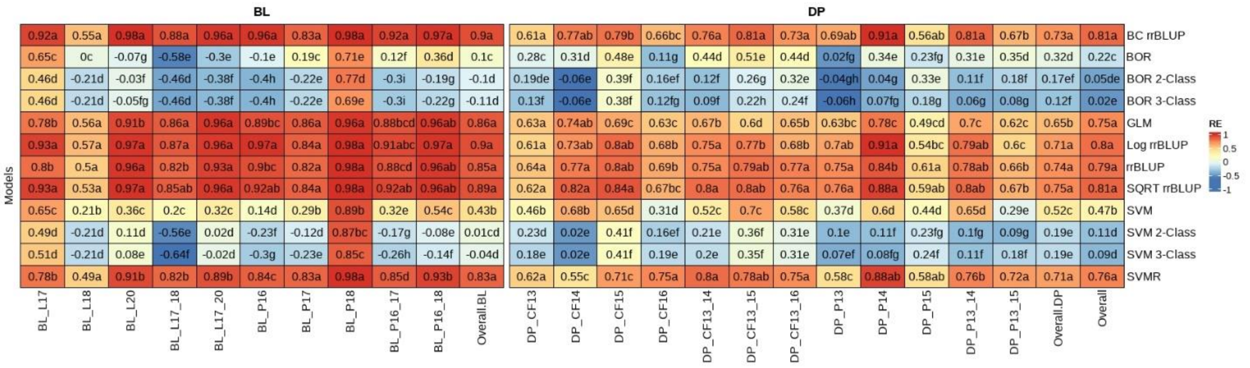


**Figure S7.** Heatmap of genomic selection regression and classification model relative efficiency (RE) and pairwise comparisons using cross-validations for stripe rust disease severity. Pacific Northwest winter wheat diversity panel (DP) lines phenotyped from 2013 to 2016 in Central Ferry (DP_CF) and Pullman (DP_P), WA. Washington State University breeding lines phenotyped from 2016 to 2020 in Lind (BL_L) and Pullman (BL_P), WA. Model comparison across the DP (Overall.DP), BL (Overall.BL), and Overall scenarios. Models labeled with the same letter are not significantly different (*P*-value =0.05).

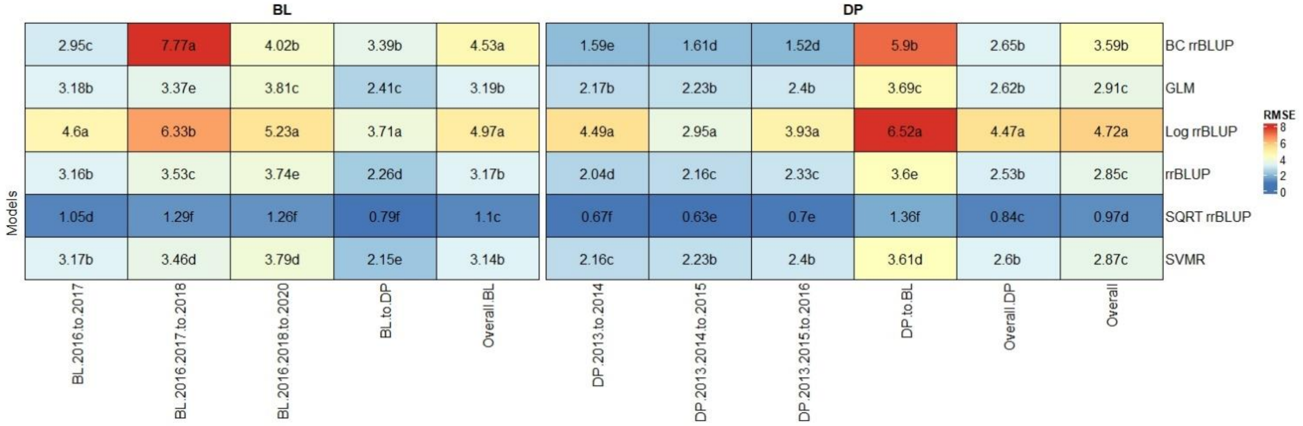


**Figure S8.** Heatmap of genomic selection regression model RMSE and pairwise comparisons using validation sets for stripe rust infection type. Pacific Northwest winter wheat diversity panel (DP) lines phenotyped from 2013 to 2016 in Central Ferry (DP_CF) and Pullman (DP_P), WA. Washington State University breeding lines phenotyped from 2016 to 2020 in Lind (BL_L) and Pullman (BL_P), WA. Model comparison across the DP (Overall.DP), BL (Overall.BL), and Overall scenarios. Models labeled with the same letter are not significantly different (*P*-value =0.05).

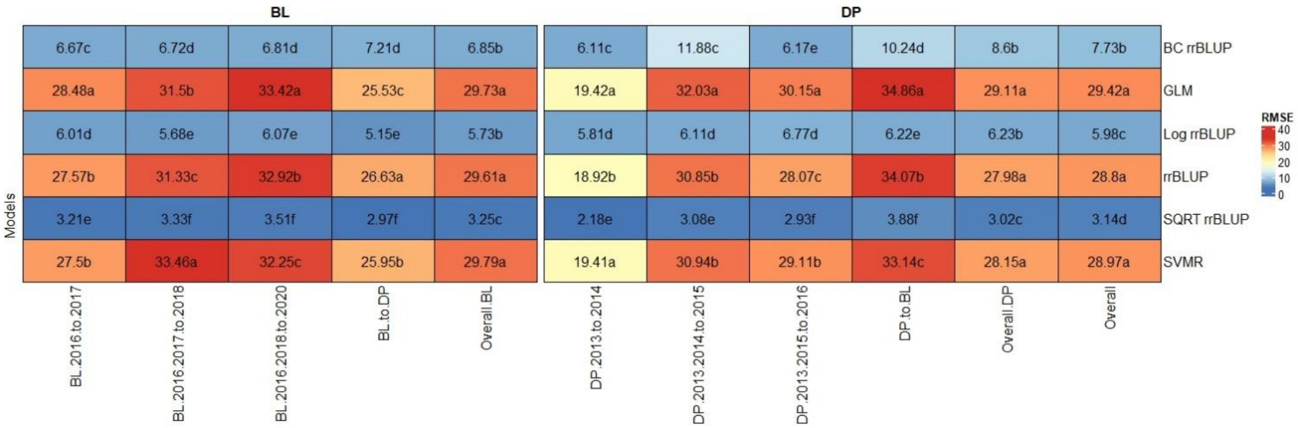


**Figure S9.** Heatmap of genomic selection regression model RMSE and pairwise comparisons using validation sets for stripe rust disease severity. Pacific Northwest winter wheat diversity panel (DP) lines phenotyped from 2013 to 2016 in Central Ferry (DP_CF) and Pullman (DP_P), WA. Washington State University breeding lines phenotyped from 2016 to 2020 in Lind (BL_L) and Pullman (BL_P), WA. Model comparison across the DP (Overall.DP), BL (Overall.BL), and Overall scenarios. Models labeled with the same letter are not significantly different (*P*-value =0.05).

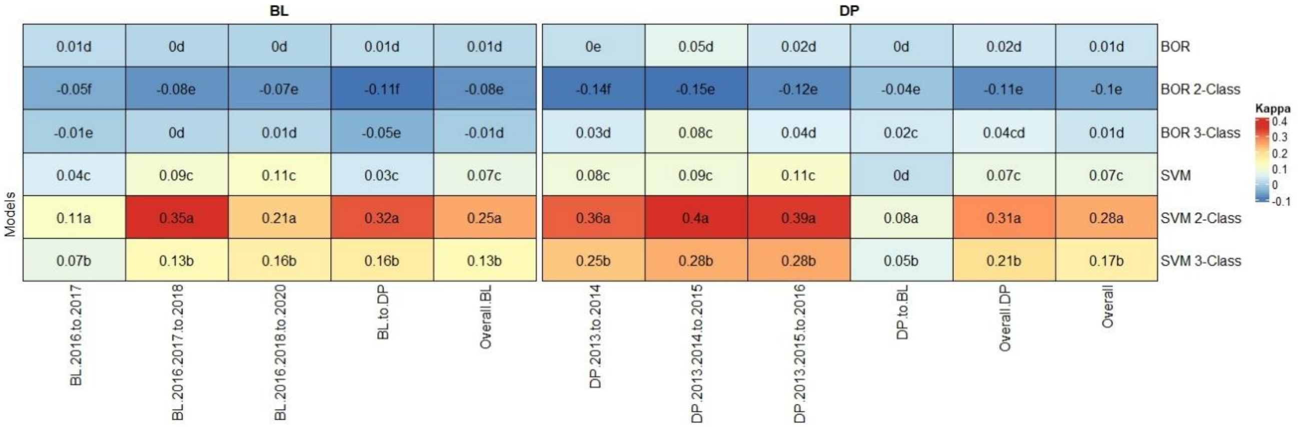


**Figure S10.** Heatmap of genomic selection classification model kappa and pairwise comparisons using validation sets for stripe rust infection type. Pacific Northwest winter wheat diversity panel (DP) lines phenotyped from 2013 to 2016 in Central Ferry (DP_CF) and Pullman (DP_P), WA. Washington State University breeding lines phenotyped from 2016 to 2020 in Lind (BL_L) and Pullman (BP_L), WA. Model comparison across the DP (Overall.DP), BL (Overall.BL), and Overall scenarios. Models labeled with the same letter are not significantly different (*P*-value =0.05).

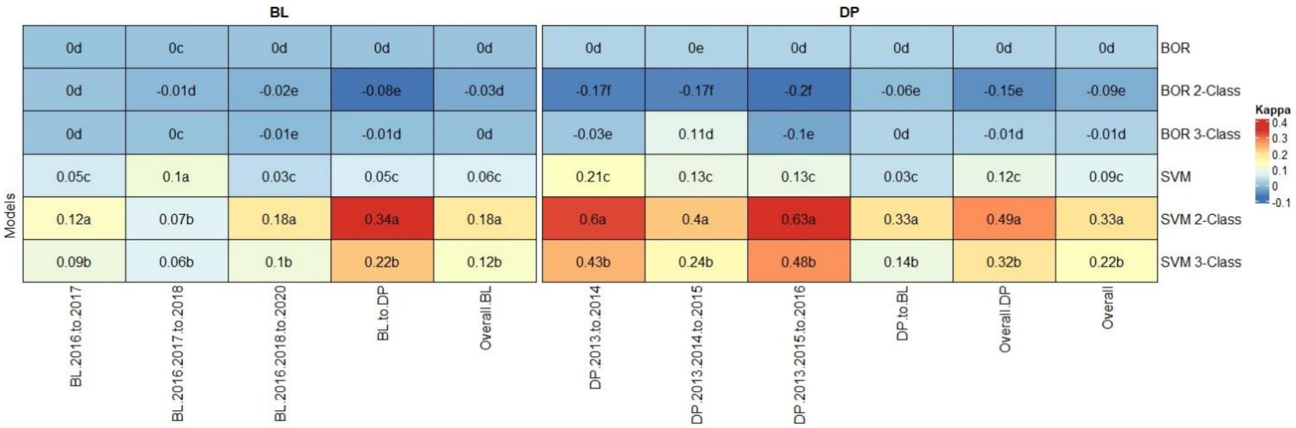


**Figure S11.** Heatmap of genomic selection classification model kappa and pairwise comparisons using validation sets for stripe rust disease severity. Pacific Northwest winter wheat diversity panel (DP) lines phenotyped from 2013 to 2016 in Central Ferry (DP_CF) and Pullman (DP_P), WA. Washington State University breeding lines phenotyped from 2016 to 2020 in Lind (BL_L) and Pullman (BL_P), WA. Model comparison across the DP (Overall.DP), BL (Overall.BL), and Overall scenarios. Models labeled with the same letter are not significantly different (*P*-value =0.05).
